# *SMAD4* loss drives cancer chromosomal instability through transcription-replication conflict-induced replication stress

**DOI:** 10.64898/2025.12.17.694873

**Authors:** Atmika Paul, Gargee Joshi, Philipp Jungk, Simranjeet Kaur, Shengyu Yao, Ioannis Tsirkas, Nicolas Böhly, Maren Sitte, Gabriela Salinas, Stephan Hamperl, Maik Kschischo, Holger Bastians

## Abstract

Cancer chromosomal instability (CIN) drives tumor evolution through generating structural and numerical chromosome aberrations. However, its molecular determinants remain poorly characterized. Here, we uncover a yet unrecognized role for the tumor suppressor *SMAD4* as a key regulator of genome stability. Mechanistically, loss of *SMAD4* induces replication stress by promoting transcription–replication conflicts (TRCs), which induces chromosomal breaks and subsequent mitotic errors, leading to the concomitant emergence of structural and numerical CIN. While the tumor suppressor function of *SMAD4* has been predominantly attributed to TGF-β signaling, we find that its role in genome maintenance operates via BMP signaling and its transcriptional target gene *ID3*. Consequently, inhibition of BMP signaling mimics the loss of *SMAD4* whereas re-expression of *SMAD4* or *ID3* suppresses TRCs, replication stress and mitotic errors in *SMAD4*-deficient cancer cells. Our findings reveal a TRC- and replication stress-driven mechanism by which loss of BMP-*SMAD4* signaling causes cancer chromosomal instability.

## Introduction

Chromosomal instability (CIN) is a major hallmark of human cancer acting as a driver for tumor evolution towards aggressive, metastatic and therapy-resistant tumor development ^1^. CIN can be broadly categorized into structural CIN (S-CIN) and numerical or whole chromosomal CIN (W-CIN) leading to generation of structural chromosome aberrations and aneuploidy respectively ^2,3^. Replication stress, a condition of slowed or stalled progression of DNA replication is considered as a key condition contributing to S-CIN in cancer, while mitotic errors associated with chromosome missegregation have been linked to W-CIN ^2–5^. Interestingly, recent studies have revealed that replication stress can have also a direct impact on mitotic fidelity ^6–9^. In fact, mild replication stress, typically detected in chromosomally unstable cancer cells, is sufficient to upregulate microtubule dynamics during mitosis. This leads to positioning defects of the mitotic spindle, and triggers whole chromosome missegregation and aneuploidy ^6–13^. Thus, replication stress contributes to both, S-CIN and W-CIN, but its molecular origins in cancer are not well understood. So far, selected oncogenes including *K-R*AS, *MYC* and *CCNE1* have been suggested as triggers for replication stress by causing cellular depletion of deoxynucleotide pools or by unscheduled activation of intragenic replication origins ^4,14,15^. In addition, collisions between the transcription and replication machineries, so-called transcription-replication conflicts (TRCs) emerged as important sources for replication stress and genome instability ^16–19^. Despite the high prevalence of TRCs, replication stress and CIN in cancer, their upstream drivers remain largely to be characterized. Here, we identify *SMAD4* as a previously unrecognized regulator of CIN.

*SMAD4* is a well-established tumor suppressor, frequently inactivated during the advanced stages of gastrointestinal malignancies, particularly in colorectal cancer (CRC) and pancreatic ductal adenocarcinoma (PDAC), coinciding with high levels of chromosomal aberrations ^20,21^. *SMAD4* inactivation, most frequently through loss of chromosome 18q, strongly contributes to late-stage malignant progression, metastasis and therapy resistance of CRC and PDAC ^20–23^. Functionally, SMAD4 acts as a central transcriptional mediator of tumor growth factor beta (TGFβ) and bone morphogenetic protein (BMP) signaling pathways ^24^. In fact, canonical TGFβ-SMAD4 signaling plays a crucial role in suppressing tumor growth through induction of cell cycle arrest, apoptosis and immune surveillance and is also linked to the epithelial-to-mesenchymal transition (EMT) and metastasis ^25,26^. The role of BMP-SMAD4 signaling in cancer has been understudied, but generally involves functions in development, differentiation, and tissue homeostasis ^24^. In this study we used an integrative approach combining bioinformatic analyses and mechanistic studies to uncover that loss of BMP-regulated *SMAD4* signaling and its target gene *ID3* results in CIN, which is triggered by transcription-replication conflict-driven replication stress and consequent mitotic errors in human cancer cells.

## Results

### Loss of the tumor suppressor *SMAD4* associated with LOH at chromosome 18q is strongly correlated with CIN in CRC and PDAC

To identify cancer-relevant lesions associated with CIN in CRC and PDAC, we conducted comprehensive bioinformatic analyses using cancer data sets from The Cancer Genome Atlas (TCGA). We assessed well-established structural complexity (SCS) and numerical complexity scores (NCS) as surrogate measures for overall S-CIN and W-CIN, respectively ^27^, and correlated these scores with chromosomal arm-level aberrations detected in CRC and PDAC. Our analysis identified the deletion of chromosome 18q as one of the most significant large scale chromosomal alteration associated with high S-CIN and high W-CIN scores in CRC and PDAC patient cohorts (Fig. 1a, b), indicating a potential CIN suppressor function of the 18q locus as suggested previously^9^. Additional correlation analyses for genes located on chromosome 18q highlighted the inactivation of the tumor suppressor *SMAD4* gene located at 18q21 as most strongly associated with S-CIN and W-CIN (Fig. 1c). Moreover, copy number loss of *SMAD4* was directly linked to high CIN scores in CRC and PDAC (Fig. 1d) suggesting that *SMAD4* might have a function in regulating CIN.

**Fig. 1:**
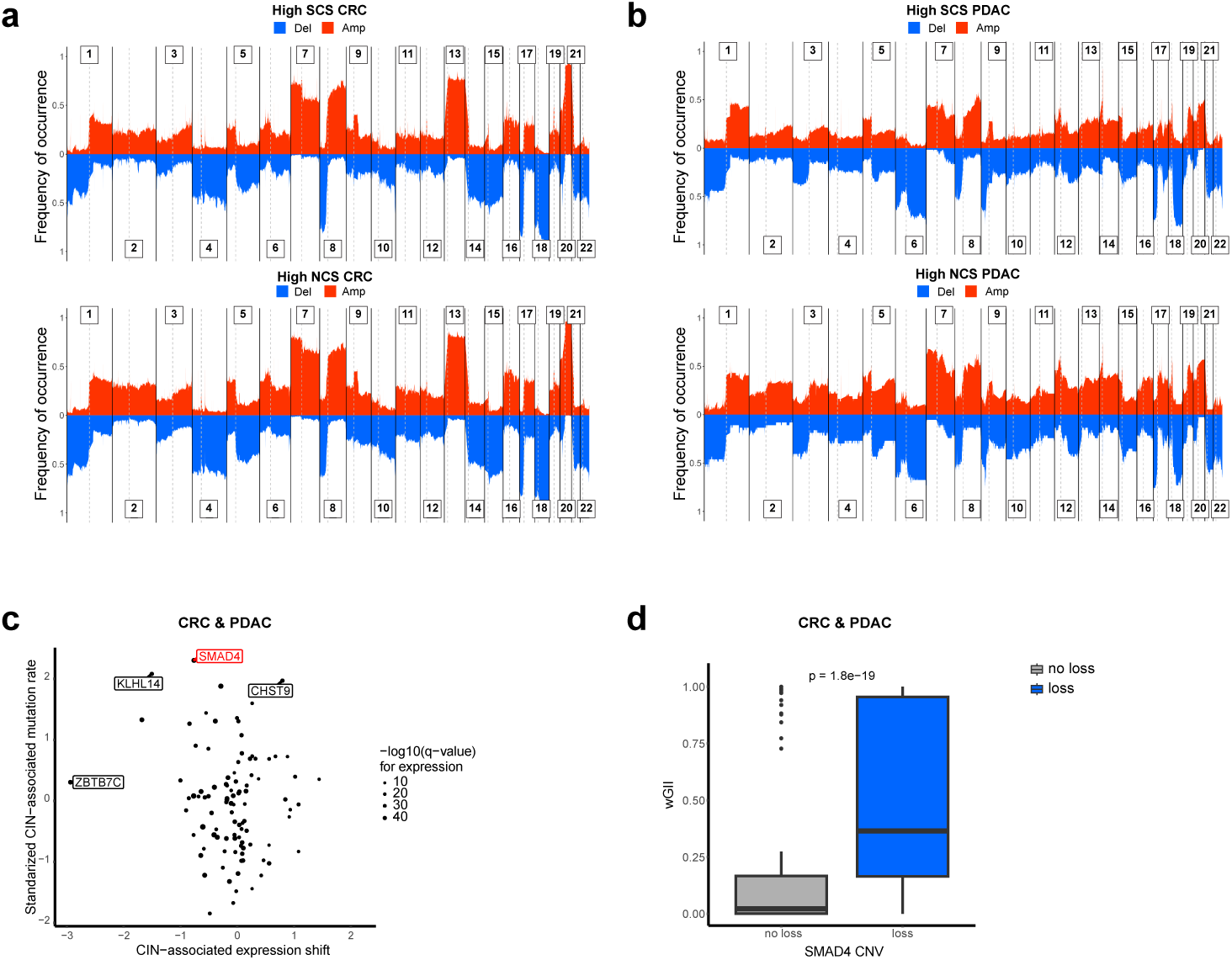
Loss of the tumor suppressor *SMAD4* on chromosome 18q is associated with structural and numerical chromosomal instability in colorectal and pancreatic cancer. **a** Association of genome-wide copy number alterations with structural (SCS) and numerical complexity scores (NCS) in colorectal cancer (CRC) (for SCS, n=239 samples; for NCS, n= 250 samples). **b** Association of genome-wide copy number alterations with structural (SCS) and numerical complexity scores (NCS) in pancreatic adenocarcinoma (PDAC) (for SCS, n=61 samples; for NCS, n= 37 samples). **c** Alterations in mRNA expression (x-axis) and mutational rates (y-axis) associated with chromosomal instability (CIN) for genes located on chromosome 18q (for expression, n=533 samples; for mutation, n=688 samples). **d** Association of *SMAD4* loss with chromosomal instability measures (wGII) in CRC and PDAC. Based on GISTIC2 copy number values, samples were classified as *SMAD4*-deficient (loss; GISTIC2 value <0) and *SMAD4*-proficient (no loss; GISTIC2 value =0; n=470 CRC and 149 PDAC samples).

### Loss of *SMAD4* triggers CIN in CRC and PDAC cells

To investigate a role of *SMAD4* in regulating CIN, we used CRISPR/Cas9-generated *SMAD4* knockout cells derived from *SMAD4*-proficient CRC and PDAC cells (Fig. 2a). Loss of *SMAD4* was not associated with growth or cell cycle impairment (Supplementary Fig. 1a). We analyzed single cell clones for the evolvement of chromosomal breaks and chromosome number variability over 30 generations as measures for S-CIN and W-CIN, respectively (Fig. 2b). We found that loss of *SMAD4* expression resulted in increased levels of chromosomal breaks as well as increased chromosome number variability in both, CRC and PDAC cells (Fig. 2c, d). Aneuploidy induction triggered by loss of *SMAD4* was further verified by CEP-FISH analysis (Supplementary Fig. 1b, left panel). In line with chromosomal breaks and aneuploidy induction, we also detected increased levels of micronuclei containing either centric (CenpC-positive), acentric (CenpC-negative) or damaged (γ−H2AX-positive) chromosomes or parts thereof, upon knockout of *SMAD4* in CRC and PDAC cells (Fig. 2e). To extend these findings to additional cellular context, we analyzed single cell clones derived from chromosomally unstable cancer cells showing endogenous loss of *SMAD4* and reconstituted *SMAD4* expression in these cells (Fig. 2f). In fact, re-expression of *SMAD4* alleviated the evolvement of chromosomal breaks, chromosome number variability, aneuploidy and micronuclei formation (Fig. 2g, h, i; Supplementary Fig. 1b, middle and right panels). Together, these results show that loss of the tumor suppressor *SMAD4* is not only associated, but sufficient to trigger chromosomal instability in human cancer cells.

**Fig. 2:**
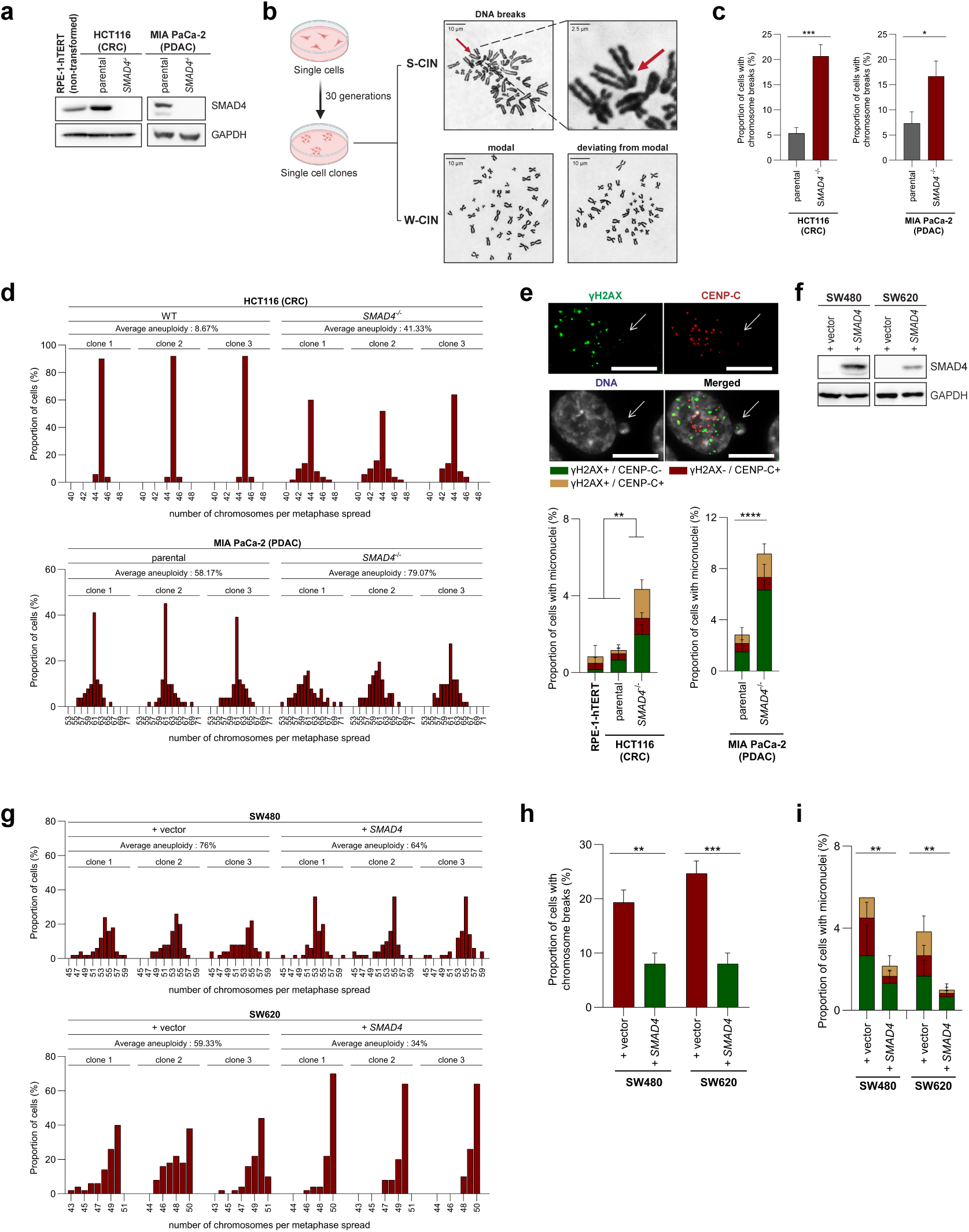
Loss of *SMAD4* triggers structural and numerical chromosomal instability. **a** Representative western blots showing loss of *SMAD4* expression in CRC (HCT116) and PDAC (MIA PaCa-2) cells with or without *SMAD4* knockout. **b** Schematic illustration of analysis of induced chromosomal breaks and numerical chromosome variability evolved in single cell clones over 30 generations. **c** Detection and quantification of chromosomal breaks in CRC (HCT116) and PDAC (MIA PaCa-2) cells with or without *SMAD4* knockout (mean ± SD, t-test, n=50 chromosome spreads per condition from 3 independent clones). **d** Quantification of chromosome number variability in CRC (HCT116) and PDAC (MIA PaCa-2) cells with or without *SMAD4* knockout and grown for 30 generations (mean ± SD, t-test, n=50 chromosome spreads per condition from 3 independent clones). **e** Micronuclei detection in non-transformed RPE-1-hTERT (control) and in CRC (HCT116) and PDAC (MIA PaCa-2) cells with or without *SMAD4* knockout using immunofluorescence microscopy detecting Cenp-C (kinetochores), γ-H2AX (DNA damage) and DNA (mean ± SD, t-test, n=600 cells from 3 experiments). Example immunofluorescence images of micronuclei are shown (scale bar, 10 µm). **f** Representative western blots showing stable re-expression of *SMAD4* in single cell clones derived from *SMAD4*-deficient SW480 and SW620 cells. **g** Quantification of chromosome number variability in *SMAD4*-deficient CRC (SW480, SW620) cells with or without stable *SMAD4* re-expression and grown for 30 generations (mean ± SD, t-test, n=50 chromosome spreads per condition from 3 independent clones). **h** Proportion of *SMAD4*-deficient cancer cells with or without *SMAD4* re-expression exhibiting chromosomal breaks (mean ± SD, t-test, n=50 chromosome spreads per condition from 3 independent clones). **i** Quantification of *SMAD4*-deficient cancer cells with or without *SMAD4* re-expression and showing micronuclei stained for Cenp-C (kinetochores), γ-H2AX (DNA damage) and DNA (mean ± SD, t-test, n=600 cells from 3 experiments).

### Loss of *SMAD4* causes CIN through transcription-replication conflicts-driven DNA replication stress

DNA replication stress is considered as a major driving force for CIN in cancer ^4^. It can be caused by various insults including nucleotide shortage, oncogene activation or transcriptional dysregulation leading to transcription-replication conflicts (TRCs) ^16,19^. RNA sequencing of CRC cells with and without *SMAD4* expression followed by gene ontology analysis revealed a strong dysregulation of pathways related to RNA polymerase II-regulated transcription, which was also reflected in *SMAD4*-deficient CRC and PDAC tumor samples (Supplementary Fig. 2a, b). This prompted us to investigate whether CIN in *SMAD4*-deficient cancer cells might be linked to TRCs and replication stress. By performing proximity-ligation-assays detecting co-localization of active RNA polymerase II and replication fork-associated PCNA ^28^, we indeed found a significant increase of TRCs in *SMAD4*-deficient CRC and PDAC cells during S-phase (Fig. 3a), while re-expression of *SMAD4* in *SMAD4*-deficient cancer cells significantly reduced the presence of TRCs (Fig. 3b) indicating that TRCs emerge due to *SMAD4* loss in cancer cells.

**Fig. 3:**
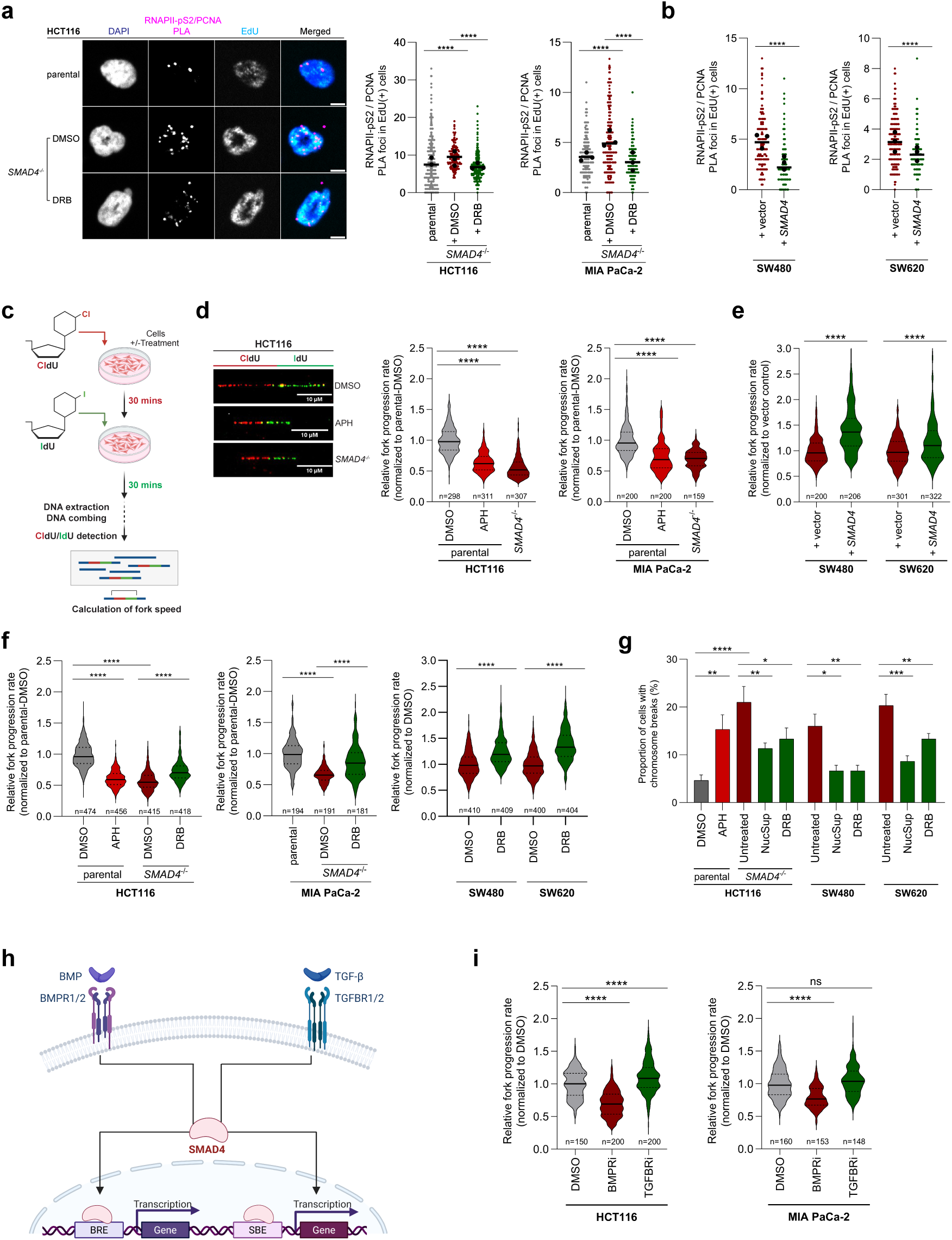
Loss of BMP-SMAD4 signaling causes TRC-mediated replication stress in cancer cells. **a** Detection and quantification of TRCs during S-phase in CRC (HCT116) and PDAC (MIA PaCa-2) cells with or without *SMAD4*-knockout. S-phase cells were labelled by EdU incorporation and TRCs were detected by PLA using RNAPII-Ser2P and PCNA antibodies. Cells were treated transiently with 100 µM DRB for 3 hours during S phase to suppress TRCs. Representative images are shown (PLA foci, magenta; DNA, blue; EdU, cyan; scale bar, 5 µm) and TRC foci were quantified. Scatter plots show individual cells from 2–3 independent biological replicates, with black dots indicating the mean of each replicate (mean, t-test, n = 427-752 cells for HCT116, 477-519 cells for MIA PaCa-2). **b** Quantification of TRC PLA foci during S-phase in *SMAD4*-deficient SW480 and SW620 cells with or without *SMAD4* re-expression. Scatter plots show individual cells from 3 independent biological replicates, with black dots indicating the mean of each replicate (mean, t-test, n = 276-378 cells for SW480, 253-665 cells for SW620). **c** Schematic illustration of DNA combing to measure velocity of individual replication forks during DNA replication. **d** Measurements of replication fork progression rates in CRC (HCT116) and PDAC (MIA PaCa-2) cells with or without *SMAD4* loss. As controls, parental cells were treated with 100 nM of aphidicolin (APH) to induce mild replication stress (normalized mean ± SD, t-test, n=159-307 DNA fibers). Example images showing CldU and IdU-labelled DNA fibers are given (scale bar, 10 µm). **e** Measurements of replication fork progression rates in *SMAD4*-deficient (SW480, SW620) cells with or without re-expression of *SMAD4* (normalized mean ± SD, t-test, n=200-322 DNA fibers). **f** Measurements of replication fork progression rates in the indicated cell lines with or without transient DRB treatment to suppress TRCs during S phase. As a control, cells were treated with 100 nM aphidicolin (APH) to induce replication stress (normalized mean ± SD, t-test, n=181-474 DNA fibers). **g** Quantification of chromosomal breaks in the indicated cell lines without or with deoxynucleoside supplementation or transient DRB treatment to alleviate replication stress and transcription-replication conflicts (TRCs), respectively. As control, parental cells were treated with 100 nM aphidicolin (APH) to induce replication stress (mean ± SD, t-test, n=50 chromosome spreads from 3 experiments). **h** Model illustrating SMAD4 as the common central mediator of the BMP and TGFβ signaling pathways. **i** Replication stress induction upon inhibition of BMP, but not TGFβ signaling. *SMAD4*-proficient CRC (HCT116) and PDAC (MIA PaCa-2) cells were treated with 1 µM BMPR inhibitor or TGFBR1/2 inhibitor and replication fork speed was measured by DNA combing (n=148-200 DNA fibers, mean ± SD, t-test).

Since TRCs might be linked to DNA replication stress, we employed DNA combing analysis as a gold-standard method to measure replication stress-induced slowing of individual replication forks (Fig. 3c) ^29^. Importantly, as seen upon direct inhibition of DNA polymerases using low concentrations of aphidicolin, loss of *SMAD4* induced replication stress in both CRC and PDAC cells (Fig. 3d) and re-expression of *SMAD4* alleviated endogenous replication stress present in *SMAD4*-deficient cancer cells (Fig. 3e). In support of this, *SMAD4*-deficient cells showed increased co-localization of FANCD2 and γ-H2AX nuclear foci during S-phase, indicative of under-replicated DNA and DNA damage, respectively ^30^ (Supplementary Fig. 2c), which was suppressed upon deoxynucleoside supplementation as an established method to alleviate replication stress ^7,9,31,32^. Also, we detected increased formation of 53BP1 nuclear bodies as another marker for replication stress ^33^ in cells treated with aphidicolin or upon loss of *SMAD4* (Supplementary Fig. 2d). Thus, loss of the tumor suppressor *SMAD4* causes TRCs and replication stress. To further investigate the link between TRCs and replication stress, we employed cell cycle synchronized cells and transiently inhibited transcription elongation during S phase using 5,6-dichloro-1-β-D-ribofuranosylbenzimidazole (DRB). This treatment did not interfere with timely progression of S phase (Supplementary Fig. 2e), but suppressed TRCs in *SMAD4*-deficient cells (Fig. 3a). Suppression of TRCs alleviated replication stress in the different *SMAD4*-deficient cancer cell systems (Fig. 3f) and also markedly suppressed the generation of chromosomal breaks similar to what is observed upon alleviation of replication stress using deoxynucleoside supplementation (Fig. 3g). Together, our results indicate that TRC-driven replication stress upon loss of *SMAD4* contributes to structural CIN.

### Loss of BMP-SMAD4, but not TGFβ-SMAD4 signaling causes replication stress

SMAD4 regulates differential gene expression in response to BMP and TGFβ signaling (Fig. 3h). Therefore, we asked whether replication stress present in *SMAD4*-deficient cells is a consequence of loss of BMP-SMAD4 or TGFβ-SMAD4 signaling. By performing DNA combing analysis, we found that inhibition of BMP, but not of TGFβ signaling is sufficient to cause replication stress in *SMAD4*-proficient CRC and PDAC cells (Fig. 3i), indicating that BMP-SMAD4, rather than TGFβ-SMAD4 signaling is required to prevent replication stress.

### *ID3* is a BMP-SMAD4 target crucial for TRC- and replication stress-associated CIN

To identify possible targets of BMP-SMAD4 signaling that are relevant to TRCs, replication stress and CIN, we performed RNA sequencing to analyze differential gene expression in distinct cell systems with or without *SMAD4* expression or with or without BMP signaling inhibition. Our integrated RNA sequencing analyses identified *inhibitor of DNA binding 3* (*ID3)*, *ID1* and *NR4A3* as genes regulated by BMP and SMAD4 signaling (Fig. 4a; Supplementary Fig. 3a). Interestingly, *ID3* has been previously characterized as a BMP-SMAD4 target gene ^34^ and implicated in the regulation of homologous DNA repair and DNA damage signaling ^35,36^. This prompted us to focus on *ID3* as a potential mediator of CIN in *SMAD4*-deficient cancer cells. First, we verified the *SMAD4*- and BMP-dependency of *ID3* mRNA and protein expression in our different CRC and PDAC cancer cell systems (Supplementary Fig. 3b, c). Importantly, the significant positive correlation between *SMAD4* and *ID3* expression was also seen upon analyzing gene expression data from CRC and PDAC tumor samples (Fig. 4b) suggesting that *ID3* expression is regulated by SMAD4 also in tumors. To investigate the role of ID3 as a SMAD4-regulated mediator in CIN-causing mechanisms, we re-expressed *ID3* in *SMAD4*-knockout CRC and PDAC cells (Supplementary Fig. 3d) and analyzed the presence of TRCs and replication stress. In fact, TRCs and replication stress present in *SMAD4*-knockout CRC and PDAC cells were both decreased upon *ID3* re-expression (Fig. 4c, e). Consistent with these results, re-expression of *ID3* in cancer cells with endogenous *SMAD4* deficiency (Supplementary Fig. 3e) also suppressed TRCs and alleviated replication stress (Fig. 4d, f) indicating that loss of *ID3* is crucial for the induction of TRCs and replication stress in *SMAD4*-deficient cancer cells. In further support of this, sole CRISPR/Cas9-mediated *ID3* knockout in *SMAD4*-proficient CRC and PDAC cells (Supplementary Fig. 3c) was sufficient to cause replication stress, similar to treatment with aphidicolin (Fig. 4e). Moreover, replication stress induced upon *ID3* knockout in CRC and PDAC cells was alleviated by transient inhibition of transcription elongation mediated by DRB treatment (Fig. 4g), which we showed to mitigate the generation of TRCs (Fig. 3a), suggesting that replication stress in both, *SMAD4*- and *ID3*-knockout cells, is TRC dependent. Finally, we investigated the generation of chromosomal breaks as a consequence of replication stress. Similar to *SMAD4* deficiency (Fig. 2c), loss of *ID3* was associated with chromosomal breaks while re-expression of *ID3* in *SMAD4*-deficient cancer cells suppressed the generation of chromosomal breaks similar to suppression of TRCs (DRB treatment) or alleviation of replication stress upon deoxynucleoside treatment (Fig. 4h). Together, our results demonstrate that loss of BMP-SMAD4 signaling, resulting in loss of *ID3* expression, is crucial for the generation of TRCs and replication stress and linked to the induction of CIN.

**Fig. 4:**
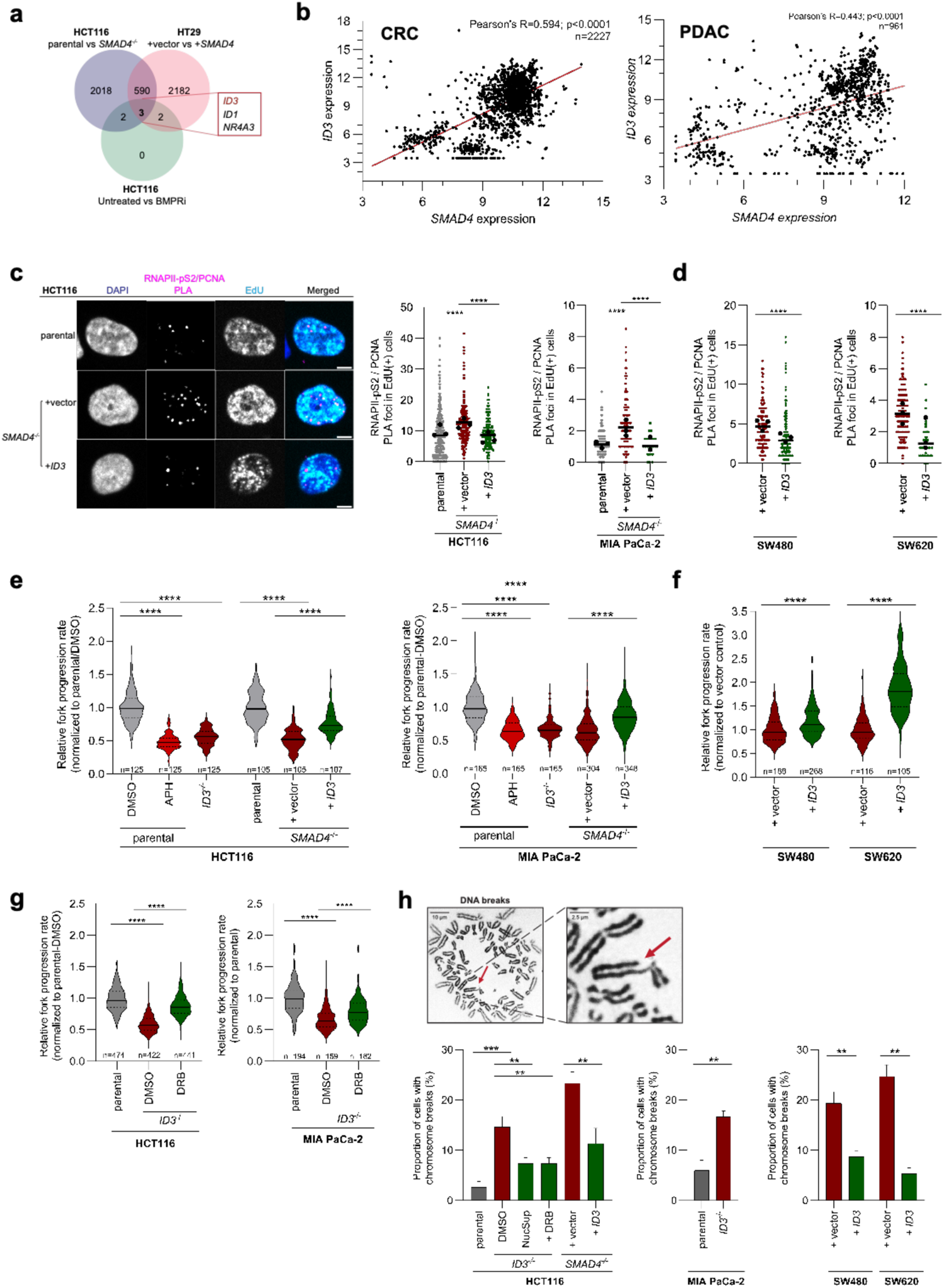
I*D*3 is a BMP-SMAD4 target crucial for TRC- and replication stress-associated CIN. **a** Venn diagrams of differentially expressed genes in HCT116 cells with or without BMP inhibition, or *SMAD4* knockout and *SMAD4*-deficient HT29 cells with or without *SMAD4* re-expression. The numbers of unique and overlapping genes are displayed. **b** Correlation analysis of *SMAD4* and *ID3* mRNA expression in CRC (n=2227) and PDAC (n=961) samples (Pearsońs correlation coefficient (R), t-test). **c** Detection and quantification of TRC PLA foci during S-phase of CRC (HCT116) and PDAC (MIA PaCa-2) *SMAD4* knockout cells with or without *ID3* re-expression. Representative images are shown (PLA foci, magenta; DNA, blue; EdU, cyan; scale bar, 5 µm). Scatter plots show individual cells from 2–3 independent biological replicates, with black dots indicating the mean of each replicate (mean, t-test, n = 271-466 cells for HCT116, 117-833 cells for MIA PaCa-2). **d** Quantification of TRC PLA foci in *SMAD4*-deficient SW480 and SW620 cells with or without *ID3* re-expression. Scatter plots show individual cells from 3 independent biological replicates, with black dots indicating the mean of each replicate (mean, t-test, n = 276-534 cells for SW480, 409-665 cells for SW620, vector control as in Fig. 3b). **e** Measurements of replication fork progression rates in CRC (HCT116) and PDAC (MIA PaCa-2) cells with or without *ID3* knockout and in *SMAD4*-knockout cells after *ID3* re-expression. As a control, cells were treated with 100 nM aphidicolin (APH) to induce replication stress (normalized mean ± SD, t-test, n=105-348 DNA fibers). **f** Measurements of replication fork progression rates in *SMAD4*-deficient SW480 and SW620 cells with or without *ID3* re-expression (normalized mean ± SD, t-test, n=105-268 DNA fibers). **g** Measurements of replication fork progression rates in HCT116- and MIA PaCa-2-*ID3* knockout cells with or without transient S-phase specific DRB treatment to alleviate TRCs (normalized mean ± SD, t-test, n=151-474 DNA fibers; parental controls as in Fig. 3f). **h** Detection and quantification of chromosomal breaks in *ID3*-knockout CRC (HCT116) and PDAC (MIA PaCa-2) cells, without or with nucleoside supplementation (NucSup) or DRB treatment to alleviate replication stress and TRCs respectively, and in *SMAD4*-deficient cells upon *ID3* re-expression (mean ± SD, t-test, n=50 chromosome spreads from 3 experiments). Representative images of chromosomal breaks are shown.

### Loss of the BMP-*SMAD4-ID3* signaling axis causes replication stress-induced mitotic errors

We demonstrated that *SMAD4* deficiency causes the generation of replication stress-mediated chromosomal breaks and also results in numerical chromosome aberrations and aneuploidy in both CRC and PDAC cells (Figs. 1, 2). In fact, recent work indicated that replication stress can affect mitosis by causing aberrantly increased spindle microtubule growth rates, leading to whole chromosome missegregation, thereby possibly explaining the concomitant presence of structural and numerical chromosome aberrations in cancer ^7–10^ (Fig. 5a). By using live cell tracking of individual microtubule plus tips, we determined mitotic microtubule growth rates in CRC and PDAC cells with or without BMP signaling, *SMAD4* or *ID3*. All conditions of inhibition of the BMP-SMAD4-ID3 axis increased mitotic microtubule growth rates to levels typically detected in chromosomally unstable cancer cells that are endogenously deficient of *SMAD4* (Fig. 5b, d; Supplementary Fig. 4a). Importantly, increased microtubule growth rates directly correlated with increased rates of chromosome missegregation (Fig. 5c, e; Supplementary Fig. 4b). Restoration of proper microtubule growth rates either upon re-expression of *SMAD4* or *ID3* or upon direct suppression of microtubule plus end growth by sub-nanomolar doses of Taxol, rescued chromosome missegregation (Fig. 5b, c, d, e; Supplementary Fig. 4a, b), verifying previous findings that increased microtubule growth rates are causally linked to whole chromosome missegregation in cancer cells ^8,12^. Finally, we investigated whether TRC-mediated replication stress seen in *SMAD4-* or *ID3*-deficient cells (Fig. 3, Fig. 4) is linked to mitotic errors. Indeed, suppression of TRCs by transient inhibition of transcription elongation or alleviation of replication stress restored proper mitotic microtubule growth rates and suppressed chromosome missegregation in CRC and PDAC cells (Fig. 5d, e, f, g; Supplementary Fig. 4c, d). Together, these findings link TRC-driven replication stress to mitotic errors, thereby driving the concomitant generation of S-CIN and W-CIN in *BMP-SMAD4-ID3*-deficient cancer cells (Fig. 6).

**Fig. 5:**
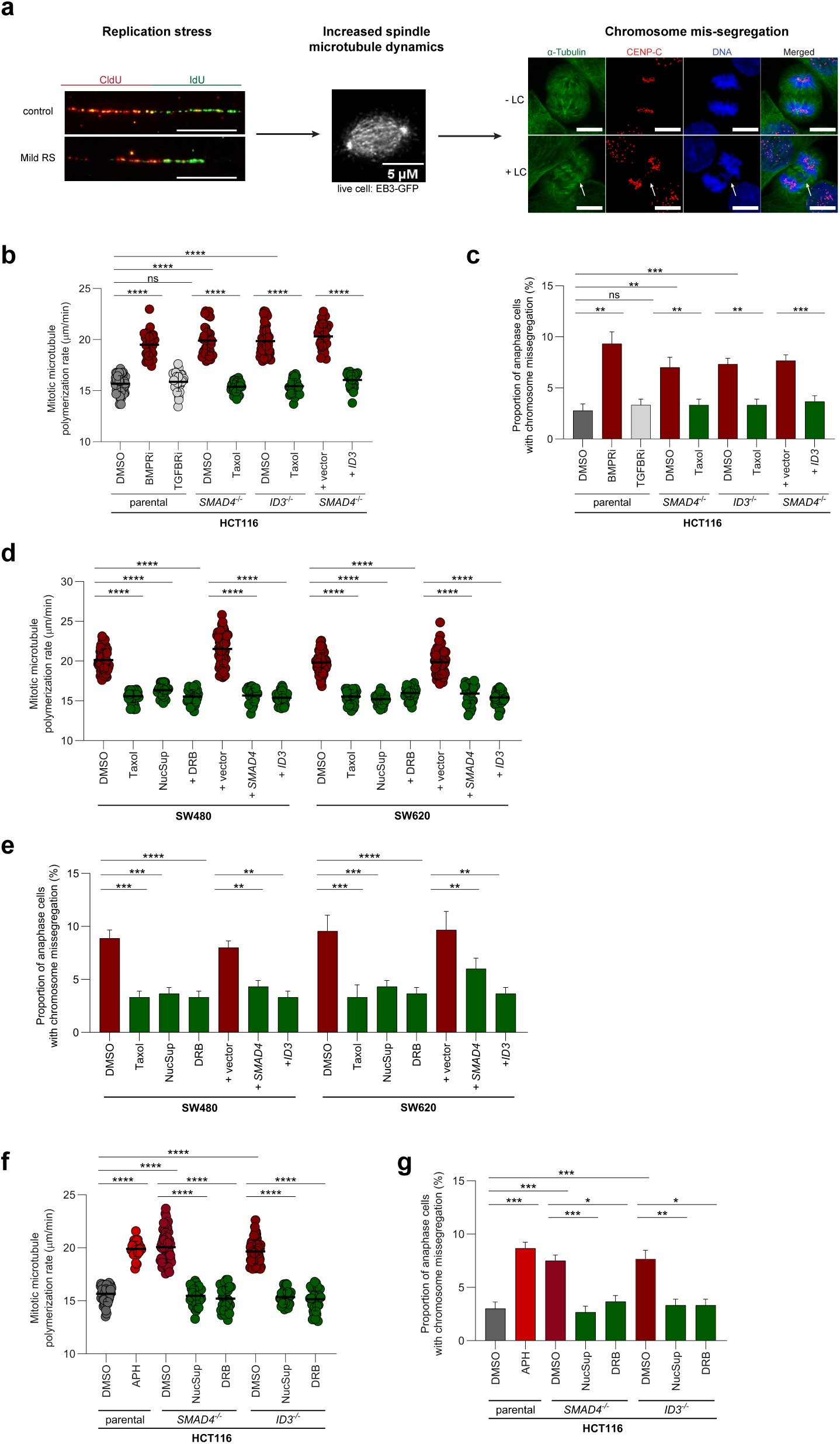
TRC- and replication stress-driven mitotic errors in *SMAD4* deficient cancer cells. **a** Schematic illustrating replication stress-induced abnormally increased mitotic microtubule growth rates leading to chromosome missegregation in mitosis. Example images show DNA fibers in the absence or presence of replication stress, an example of a mitotic cell expressing EB3-GFP to track individual microtubule plus tips in living cells and examples of cells with or without chromosome missegregation, as indicated by the presence of lagging chromatids during anaphase. **b** Measurements of mitotic microtubule growth rates in HCT116 cells treated with or without BMP or TGFβ receptor inhibition, and with and without *SMAD4-* or *ID3-* knockout, and with or without *ID3* re-expression in *SMAD4*-deficient cells. Cells were treated with 0.2 nM Taxol to correct abnormally increased microtubule growth rates (mean ± SD, t-test, n=30 cells from 3 experiments with total of 600 microtubules measured). **c** Quantification of cells showing chromosome missegregation using cells treated as in (b) (mean ± SD, t-test, n=300 anaphase cells from 3 experiments). **d** Measurements of mitotic microtubule growth rates in *SMAD4*-deficient SW480 and SW620 cells, without or with nucleoside supplementation (NucSup) or DRB treatment to alleviate replication stress and TRCs respectively, and with and without *SMAD4* or *ID3* re-expression. Cells were treated with 0.2 nM Taxol to correct abnormally increased microtubule growth rates (mean ± SD, t-test, n=30 cells from 3 experiments with total of 600 microtubules measured. **e** Quantification of cells showing chromosome missegregation using cells as in (d) (mean ± SD, t-test, n=300 anaphase cells from 3 experiments). **f** Measurements of mitotic microtubule growth rates in *SMAD4*- or *ID3*- knockout cells, without or with nucleoside supplementation (NucSup) or DRB treatment. Cells were treated with 100 nM aphidicolin (APH) to induce replication stress (mean ± SD, t-test, n=30 cells from 3 experiments with total of 600 microtubules measured). **g** Quantification of cells showing chromosome missegregation using cells treated as in (f) (mean ± SD, t-test, n=300 anaphase cells from 3 experiments).

**Fig. 6:**
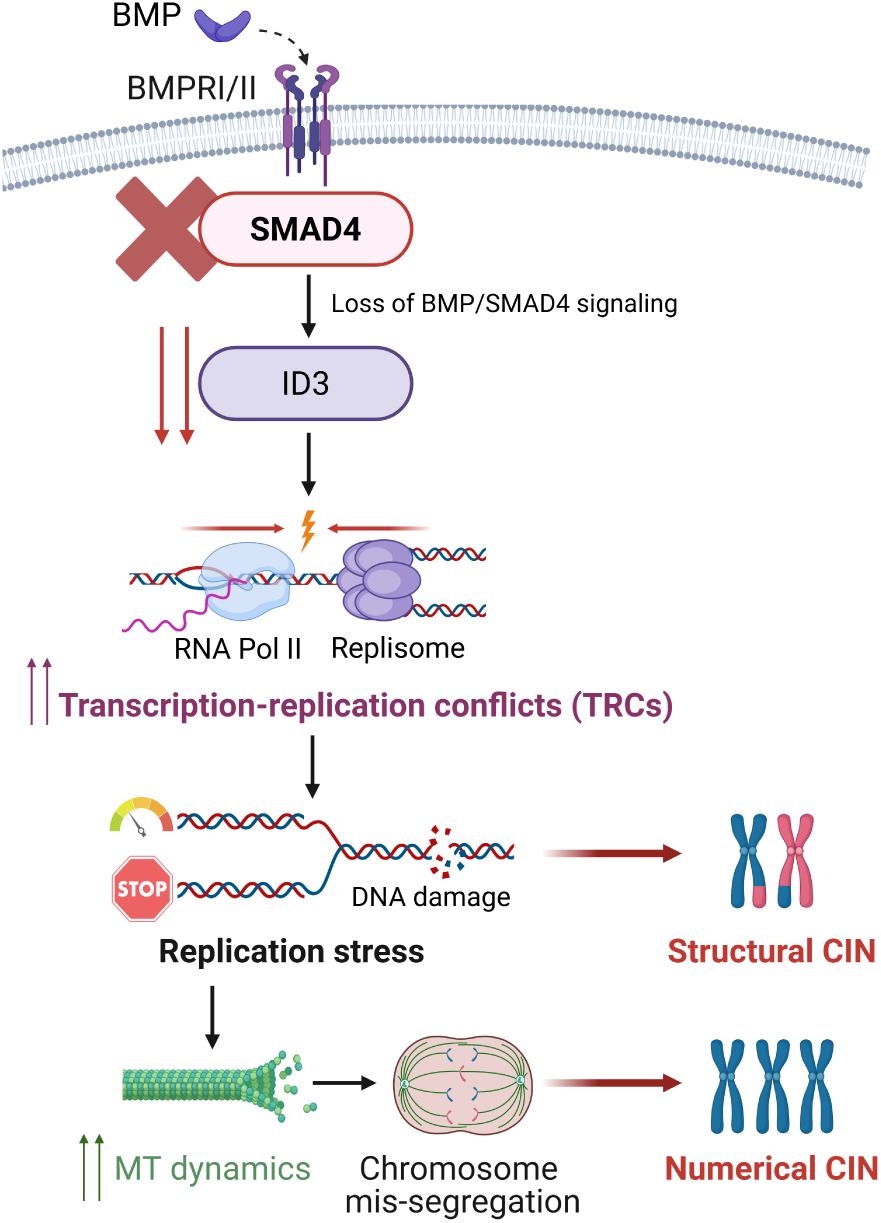
Model summarizing the role of BMP-SMAD4-ID3 signaling in promoting transcription-replication conflict-driven replication stress and mitotic errors, leading to cancer chromosomal instability (CIN).

## Discussion

*SMAD4* is one of the most frequently inactivated tumor suppressor genes in human cancer, particularly in CRC and PDAC. It is typically inactivated during late tumorigenesis and strongly supports malignant tumor progression and therapy resistance via poorly understood mechanisms ^20,22,37,38^. In addition to inactivating mutations, LOH of chromosome 18q where *SMAD4* is localized, is considered as a major mechanism of *SMAD4* inactivation ^9,23,39^. Our study showed that loss of chromosome 18q associated with reduced *SMAD4* expression is most significantly linked to chromosomal instability (CIN) in CRC and PDAC tumor samples. Interestingly, a previous study suggested that additional genes located on chromosome 18q including *PIGN*, *MEX3C* and *ZNF516*, which have not been further functionally characterized, are also associated with CIN suggesting that chromosome 18q might have a broader CIN suppressor activity ^9^. However, we demonstrate that the sole loss of *SMAD4* is sufficient to trigger CIN, both on a structural and numerical chromosome level. This is mediated by triggering transcription-replication conflicts (TRCs) during DNA replication causing replication stress and subsequent mitotic errors. Thus, our findings provide strong evidence for a potential, but yet unrecognized tumor suppressor function of *SMAD4* in maintaining genome stability, which is highly relevant during late stages of tumor progression when *SMAD4* is typically inactivated.

So far, SMAD4 has been mainly implicated as a transcription factor acting downstream of TGFβ signalling and involved in impacting cell cycle progression and apoptosis as well as in mediating epithelial-to-mesenchymal transition (EMT) in cancer ^25,26^. In contrast to that, we find that BMP-SMAD4, rather than TGFβ-SMAD4 signaling is relevant for its role in CIN suppression. Moreover, we identified *ID3* as a BMP-SMAD4 target gene that suppresses TRCs, replication stress and mitotic errors, and consequently, CIN. Notably, loss of *SMAD4* correlates with low *ID3* expression in CRC and PDAC tumor samples and low *ID3* expression has been associated with tumor progression, metastasis and poor survival outcomes ^40^, all of which typically associated with high CIN. Interestingly, ID3 belongs to the family of “inhibitor of DNA binding (ID)” proteins that act as dominant negative regulators of basic helix-loop-helix transcription factors ^41^. Hence, one could speculate that hyper-activity of ID3-regulated transcription factors as a result of *SMAD4* deficiency could lead to local hyper-transcription that might be linked to transcription-replication conflicts as a cause for replication stress. Our RNA sequencing analysis of *SMAD4*-deficient cells support this view by showing broad deregulation of RNA polymerase II-regulated gene transcription upon loss of *SMAD4*, both in cell lines and in tumor samples. Consequently, transient inhibition of transcriptional elongation suppressed both TRCs and replication stress in *SMAD4*-and *ID3*-deficient cells. In this regard, it is interesting to note that oncogene-induced hyper-transcription has also been linked to replication stress in cancer, but the mechanisms of how oncogene activation induce TRCs remain unclear ^42^. On the other hand, ID3 has been also implicated in DNA damage responses and repair ^35,36^ and thus, it is possible that impairment of these functions might be involved in increasing TRCs and replication stress. An example for such a scenario is the inactivation of regulators of homologous recombination repair such as *BRCA1* or *RAD51*, which cause not only impairment of HR repair, but also increase TRCs and slow-down of replication fork progression ^17,31^.

Typically, chromosomally unstable cancer cells are characterized by the concomitant presence of structural and numerical chromosome aberrations. This might be explained by recent studies showing that replication stress is not only associated with DNA damage due to e.g. replication fork instability and fork collapse, but also affects the faithful progression of the subsequent mitosis ^6–11^. In fact, both experimentally induced as well as endogenous replication stress in chromosomally unstable cancer cells have been shown to cause an abnormal increase in spindle microtubule dynamics. This affects correct spindle positioning, thereby leading to whole chromosome missegregation and aneuploidy evolvement ^6–10^. In this study, we found that *SMAD4* and *ID3*-deficient cancer cells exhibit mitotic errors, which are due to increased microtubule growth rates and triggered by TRC-mediated replication stress. Hence, replication stress and mitotic errors, which are commonly detected in chromosomally unstable cancer cells, appear to be frequently caused by loss of *BMP*-*SMAD4-ID3* signaling, making loss of *SMAD4*, in particular, a key cancer-associated lesion involved in CIN.

It is assumed that high CIN is present during late stages of tumor progression, which coincides with the loss of chromosome 18q and loss of *SMAD4* expression, thereby further supporting the link between loss of *SMAD4* and CIN ^23^. However, the generation of chromosomal aberrations is not restricted to late stage tumors, but can also appear early during tumor progression raising the question regarding the trigger of an early CIN scenario. Our work revealed that inhibition of BMP signaling is sufficient to trigger TRC-driven replication stress and also subsequent mitotic errors, even in the absence of *SMAD4* inactivation, indicating that the mere, possibly transient, inhibition of BMP signaling, either on a BMP ligand or BMP receptor level, is sufficient to cause CIN, which might be relevant already in early cancer stages. In fact, patients suffering from *familial juvenile polyposis syndrome*, an autosomal dominant disease with predisposition to the development of gastrointestinal cancers, show mutations in *BMPR1A* coding for a BMP receptor indicating that inhibition of BMP signaling is present in pre-cancerous lesions ^43^. Along the same lines, recent reports showed that BMP signals can protect mouse embryonic stem cells and zebrafish cardiomyocytes from replication stress during pluripotency and heart regeneration, respectively, supporting a link between BMP signaling, replication stress and CIN ^44,45^. Thus, it is tempting to speculate that modulating BMP signaling, in the absence of any stable genetic lesion in the pathway, might provide a basis for transient occurrence of CIN during early tumorigenesis, which might be later on manifested through subsequent selection for stable genetic lesions such as *SMAD4* deletion.

## Methods

### Cell culture

HCT116, SW480, SW620, HT29 cells and their *SMAD4*- and *ID3*-knockout or *SMAD4/ID3* re-expression derivatives were cultured in RPMI-1640 medium (Anprotec, Germany). MIA PaCa-2 cells and derivatives were maintained in DMEM (PAN-Biotech GmbH, Germany). All cancer cell lines were obtained from ATCC (Manassas, USA) and tested for authenticity. Non-transformed human retinal pigment epithelial (RPE-1-hTERT) cells were kindly provided by Zuzana Storchova (Kaiserslautern, Germany) and were cultured in DMEM-F12 medium supplemented with 0.26% NaHCO₃. All media were supplemented with 10% fetal bovine serum (FBS; Anprotec, Germany), 100 units/ml penicillin, and 100 µg/ml streptomycin (Anprotec, Germany). Cells were incubated at 37°C in a humidified atmosphere with 5% CO₂.

### Cell treatments

To suppress transcription-replication conflicts (TRCs), cells were treated transiently during S phase with 100 µM of the RNA Polymerase II inhibitor 5,6-dichloro-1-β-D-ribofuranosylbenzimidazole (DRB; Focus Biomolecules, USA). To induce replication stress, cells were treated with 100–200 nM aphidicolin (Santa Cruz Biotechnology, USA). For inhibition of BMP signaling, cells were treated with 1 µM Type I BMP receptor inhibitor (K02288, Selleckchem, USA). Inhibition of the TGFβ receptor I/II was achieved using 0.5–1 µM LY2109761 (Selleckchem, USA). To restore normal microtubule polymerization rates, cells were treated with 0.2 nM Taxol (Sigma-Aldrich, Germany).

### Plasmids and transfection

For *SMAD4* expression, pcDNA3-FLAG-SMAD4 was kindly provided by Aristidis Moustakas (Addgene #80888; ^46^). To express *ID3*, the pcDNA3.1+/-ID3-C-(K)-DYK plasmid was purchased from Genscript (Netherlands). The pcDNA3.1-empty vector control was purchased from Invitrogen (USA). Additionally, the pLVX-ID3-GFP plasmid and pLVX-empty (or pLVX-Puro) were both kindly provided by Ali Bakr (Heidelberg, Germany; ^35^). The pEGFP-EB3 plasmid, encoding GFP-tagged EB3 for visualizing microtubule +TIPs, was generously provided by Linda Wordeman (Seattle, WA, USA). Plasmid DNA was transfected into cells using either electroporation with a GenePulser Xcell (Bio-Rad Laboratories, USA) at 500 μF and 300 V or lipid-based transfection with Lipofectamine 3000 (Thermo Fisher Scientific, USA) according to the manufacturer’s protocol. Ectopic protein expression was confirmed by Western blot analysis.

### Generation of knockout cell lines

HCT116-*SMAD4* knockout cells were purchased from Horizon Discovery Group plc, UK. HCT116-*ID3^-/-^,* MIA PaCa-2*-SMAD4^-/-^* and MIA PaCa-2*-ID3^-/-^* cell lines were generated using the CRISPR-Cas9 gene editing system. Guide RNA (gRNA) sequences (Supplementary Table 1) targeting exon regions of the *SMAD4* and *ID3* open reading frames were designed using the IDT Alt-R® CRISPR-Cas9 gRNA designer tool (Integrated DNA Technologies, USA). The crRNA oligonucleotides were synthesized as Alt-R® CRISPR-Cas9 crRNA (Integrated DNA Technologies, USA) and combined with tracrRNA oligos (Integrated DNA Technologies, USA) in nuclease-free duplex buffer (Integrated DNA Technologies, USA) to form 1 μM crRNA:tracrRNA duplexes. Annealing was performed by heating the mixture to 95°C for 5 minutes, followed by cooling to room temperature (25°C). To form the ribonucleoprotein (RNP) complex, 1 μM of Alt-R™ Streptococcus pyogenes HiFi Cas9 Nuclease V3 (Integrated DNA Technologies, USA) was mixed with the crRNA:tracrRNA duplex and incubated at room temperature for 5 minutes. The RNP complexes were then transfected into cells using the Alt-R™ CRISPR-Cas9 System for cationic lipid-mediated transfection (Integrated DNA Technologies, USA) according to the manufacturer’s instructions. Forty-eight hours post-transfection, cells were seeded into 96-well plates for single-cell cloning. Individual clones were expanded for 30 generations, and knockout efficiency for each clone was validated through Sanger sequencing, qRT-PCR, and Western blot analysis. Clones confirmed as knockouts were subsequently used for further analysis. The primer sequences used for target amplification and Sanger sequencing are listed in Supplementary Table 2.

### Generation of transgene-expressing cell lines

To generate cell lines stably expressing *SMAD4* or *ID3*, SW480, SW620 and HT29 cells were transfected with 2.5 µg of either pcDNA3-FLAG-SMAD4, pcDNA3.1+/-ID3-C-(K)-DYK, or pcDNA3.1-empty vector using Lipofectamine 3000 (Thermo Fisher Scientific, USA) according to the manufacturer’s protocol. Single cell clones were grown, isolated and cultured in 300 μg/ml G418 selection medium for 30 generations. Stable expression of the transgenes was confirmed by western blotting before further analysis. To generate HCT116-*SMAD4*^-/-^ cell lines with stable re-expression of *ID3* or empty vector controls, cells were transfected with 3 µg of either pLVX-ID3-GFP or pLVX-empty vector using Lipofectamine 3000. Single-cell clones were selected in 1 μg/ml puromycin-containing medium for 30 generations, and stable expression of *ID3* was confirmed by western blotting. To generate MIA PaCa-2**-***SMAD4*^-/-^ cell lines with stable re-expression of *ID3* or empty-vector controls, cells were transfected with 2.5 µg of either pcDNA3.1(+/–)-ID3-C-(K)-DYK or the pcDNA3.1 empty vector using Lipofectamine 3000. Transfected cells were selected in 300 µg/ml G418, and stable *ID3* expression was verified by western blotting prior to further experiments.

### RNA isolation, cDNA synthesis, and qRT-PCR

Total RNA was isolated from cells using TRIzol™ reagent (Invitrogen, Carlsbad, USA) according to the manufacturer’s instructions. RNA concentration was determined using a Nanodrop™ 2000 spectrophotometer. For cDNA synthesis, 500 μg of RNA was reverse transcribed using the FastGene® Scriptase II cDNA Kit (#LS53; Nippon Genetics Europe) following the manufacturer’s protocol. The resulting cDNA was amplified using gene-specific primers (Supplementary Table 3), and qRT-PCR was performed with the qRT-PCR 2x qPCRBIO SyGreen Mix Lo-ROX (#PB20.11; PCR Biosystems), according to the manufacturer’s guidelines. mRNA expression levels of the target genes were normalized to *GAPDH*, and relative expression was calculated using the 2-ΔΔCT method ^47^.

### RNA sequencing

RNA sequencing was performed on HCT116 cells with and without *SMAD4* knockout, HCT116 cells treated with or without treatment with 1 µM Type I BMP receptor inhibitor (K02288, Selleckchem, USA) and on *SMAD4*-deficient HT29 cells with and without stable *SMAD4*-re-expression. Total RNA was isolated from cells using TRIzol™ reagent (Invitrogen, Carlsbad, USA) according to the manufacturer’s instructions. RNA quality was assessed by measuring the RNA integrity number (RIN) using a Fragment Analyzer HS Total RNA Kit (DNF-472-FR; Agilent Technologies). Library preparation for RNA sequencing was performed in the STAR Hamilton NGS automation using the Illumina Stranded mRNA Prep (Cat. N° 20040529) and the ID for Illumina RNA UD Indexes Set A, Ligation with 96 Indexes (Cat. N° 20091655) starting from 300 ng of total RNA. The size range of the final cDNA libraries was determined by applying the SS NGS Fragment 1- to 6000-bp Kit on the Fragment Analyzer (average 290-320 bp). Accurate quantification of cDNA libraries was performed by using the DeNovix DS-Series System. cDNA libraries were sequenced by using the HiSeq4000; 100 cycles, 30 Mio reads/sample from Illumina. Sequence images were transformed with BaseCaller Illumina software to BCL files and demultiplexed to fastq files with bcl2fastq v2.20.0.422. Sequencing quality was determined using FastQC v. 0.11.5 software (http://www.bioinformatics.babraham.ac.uk/projects/fastqc/). Sequence images were transformed with Illumina software BaseCaller to BCL files, which were demultiplexed to fastq files with bcl2fastq v2.20. The quality of the sequencing data was assessed using FastQC (http://www.bioinformatics.babraham.ac.uk/projects/fastqc/).

Sequences were aligned to the reference genome *Homo sapiens* (GRCh38.p13, https://www.ensembl.org/ Homo_sapiens/Info/Index) using the RNA-Seq alignment tool STAR version 2.7.8a ^48^. The alignment was performed with a tolerance of 2 mismatches within 50 bases. Subsequently, read counting was performed using featureCounts ^49^. The resulting read counts were analyzed in the R/Bioconductor environment (version 4.1.3, www.bioconductor.org) using the DESeq2 package version 1.32.0 ^50^. Candidate genes were filtered based on an absolute log2 fold-change >1 and FDR-corrected p-value <0.05.

### Western blotting

Cells were lysed in freshly prepared lysis buffer (20 mM Tris-HCl pH 7.5, 150 mM NaCl, 10 mM EDTA, 1% Triton X-100, 0.1% SDS, 1% sodium deoxycholate, 2 M urea) supplemented with PhosSTOP™ phosphatase inhibitor (Roche, Switzerland) and cOmplete™ EDTA-free protease inhibitor cocktail (Roche, Switzerland). Lysates were sonicated and centrifuged at 14,800 rpm for 15 minutes at 4°C. Protein concentration was determined using the Bio-Rad DC™ Protein Assay Kit (Bio-Rad, USA) following the manufacturer’s protocol. Proteins were resolved on 10-15% SDS-PAGE gels and transferred onto nitrocellulose membranes. Membranes were blocked with 5% non-fat milk in TBS (Tris-buffered saline) and incubated overnight at 4°C with the following primary antibodies at the indicated dilutions: anti-ID3 (1:750; D16D10; Cell Signaling, USA; cat no #9837; RRID: AB_2732885), anti-SMAD4 (1:1000; D3R4N; Cell Signaling, USA; cat no #46535S; RRID: AB_2736998) and anti-GAPDH (1:1000; G-9; Santa Cruz, USA; cat no sc-365062; RRID: AB_10847862). After primary antibody incubation, membranes were washed with TBS-T (TBS containing 0.1% Tween-20) and incubated for 1 hour at room temperature with horseradish peroxidase-conjugated secondary antibodies (1:10000; Jackson ImmunoResearch Laboratories, Inc., USA; cat no 115-035-146 and 111-035-144; RRID: AB_2307392 and AB_2307391) in 3% non-fat milk in TBS. Protein detection was performed using enhanced chemiluminescence.

### DNA combing

To assess replication stress, replication fork progression was analyzed by DNA combing. Cells were sequentially pulse-labeled with 100 μM CldU and IdU (Sigma-Aldrich, Germany) for 30 minutes each. Genomic DNA was extracted from 175,000 cells using the FiberPrep® kit (Genomic Vision, France) as previously described ^8^. Extracted DNA was combed onto vinyl silane-treated engraved coverslips (Combicoverslips; Genomic Vision, France or PolyAn’s Combing Coverslip; PolyAn, Germany) using the FiberComb® system (Genomic Vision, France) at 300 μm/sec (2 kb/μm stretching factor). Combed DNA was denatured (0.5 M NaOH, 1 M NaCl in H_2_O) at 8 min in room temperature and then blocked with BlockAid™ (Thermo Fisher Scientific, USA). Coverslips were then incubated with the following primary antibodies at the indicated dilutions: anti-BrdU (for CldU detection; 1:10; BU1/75 (ICR1), Abcam, UK, cat. no. ab6326; RRID: AB_305426), anti-BrdU (for IdU detection; 1:10; B44, BD Biosciences, USA, cat. no. 347580; RRID: AB_400326), and anti-ssDNA (1:5; DSHB, USA; RRID: AB_10805144;) and with secondary antibodies conjugated to Cy5 (1:25; Abcam, UK; cat no ab6565; RRID: AB_955063), Cy3.5 (1:25; Abcam, UK; cat no ab6946; RRID: AB_955045), Alexa 488 (1:25; Thermo Fisher Scientific, USA; cat no A-21121), Alexa 594 (1:25; Abcam, UK; cat no ab150160), BV480 (1:25; BD Biosciences, USA; cat no 564877; RRID: AB_2738995) or BV421 (1:25; BD Biosciences, USA; cat. no. 563846; RRID: AB_2738449). All antibody incubations were performed for 1 h each at 37°C in a humidified chamber. Unidirectional DNA tracks were visualized using either Genomic Vision’s EasyScan service (images acquired by Genomic Vision, France and analyzed using FiberStudio® software) or by acquiring images using a Leica DMI6000B fluorescence microscope equipped with a DFC360 FX camera (Leica Microsystems, Germany) or on a Nikon Eclipse Ti2 microscope equipped with a Hamamatsu ORCA-Flash4.0 camera (Hamamatsu Photonics, Japan) and analyzed using Leica LAS AF or NIS-Elements software.

### Deoxynucleoside supplementation

To alleviate replication stress, cells were treated with 20 µM of the following deoxynucleosides: 2ʹ-deoxyadenosine monohydrate, 2ʹ-deoxycytidine hydrochloride, thymidine, and 2ʹ-deoxyguanosine monohydrate (Santa Cruz, USA) for 48 hours, as described ^8,9,31,32^.

### Live cell microscopy to measure microtubule growth rates

To measure microtubule growth rates, EB3-GFP tracking experiments were conducted as described previously ^8,12^. Cells were transfected with the pEGFP-EB3 plasmid and 48 hours post-transfection, cells were accumulated in prometaphase by treatment with 2 μM Dimethylenastron (Sigma-Aldrich, Germany) for 1–2 hours to induce monopolar spindle formation and to ensure that growth rate measurements were conducted in the same mitotic phase as shown in ^12^. Live-cell imaging was performed using a DeltaVision ELITE microscope (GE Healthcare, UK) equipped with a PCO Edge sCMOS camera (PCO, Germany). Cells were maintained at 37°C with 5% CO₂ during imaging. Time-lapse images were captured every 2 seconds for 30 seconds, acquiring four z-stacks with 0.4 μm spacing between planes. Deconvolved images were analyzed using softWoRx® 6.0 software (GE Healthcare, UK) to calculate microtubule growth rates based on EB3-GFP track movement. For each condition, 20 microtubules per cell were analyzed in 10 mitotic cells, with experiments conducted in biological triplicates. Growth rates from 30 mitotic cells were averaged and plotted.

### Analysis of mitotic chromosome missegregation

To assess chromosome missegregation, lagging chromosomes were detected in anaphase cells. Cells were synchronized at the G1/S phase using a double thymidine block and released for 8.5–9 hours to enrich cells in anaphase as described. Cells were fixed with 2% paraformaldehyde (PFA) for 5 minutes at room temperature, permeabilized with ice-cold methanol at -20°C for 5 minutes, and washed with PBS. Blocking was performed with 5% FBS in PBS for 30 minutes at room temperature. Cells were incubated overnight at 4°C with anti-CENP-C (1:750; MBL International, USA; PD030; RRID: AB_10693556) and anti-α-tubulin (1:750; B-5-1-2, Santa Cruz Biotechnology, USA; sc-23948; RRID: AB_628410) to label centromeres and microtubule spindles. After washing, cells were incubated with Alexa Fluor®-conjugated secondary antibodies (Alexa Fluor 488 and Alexa Fluor 594; 1:1000; Thermo Fisher Scientific, USA; A-11029 and A-11076) for 1 hour at room temperature in the dark. Chromosomes/DNA were counterstained with Hoechst 33342 (1:10,000; Thermo Fisher Scientific, USA) for 5 minutes. Coverslips were mounted using VECTASHIELD® mounting medium (Vector Laboratories, USA) and analyzed using a Leica DMI6000B fluorescence microscope with a DFC360 FX camera (Leica Microsystems, Germany). Kinetochore-positive (Cenp-C-positive) lagging chromosomes in anaphase cells were quantified in 100 anaphase cells per replicate.

### Flow cytometry for cell cycle analysis

Cells were harvested and fixed by dropwise addition of 70% chilled ethanol to cell suspensions. After fixation, ethanol was removed by a single wash with PBS. The cells were treated with RNase (1 mg/ml in PBS; AppliChem GmbH, Germany) for 30 minutes at room temperature, followed by staining of cellular DNA with propidium iodide (1 μg/ml in PBS). Cell cycle profiles were analyzed by acquiring data from at least 10,000 cells per experimental condition using a BD FACSCanto™ II flow cytometer (Becton Dickinson, Germany). Data analysis was performed using BD FACSDiva™ software (Becton Dickinson, Germany) or Floreada.io (https://floreada.io/).

### Immunofluorescence microscopy and micronuclei detection

To detect S-phase cells, cells were labeled with 20 μM EdU (Jena Biosciences, Germany) for 30 minutes and EdU incorporation was detected using the Click-iT™ EdU Cell Proliferation Kit for Imaging with Alexa Fluor™ 488 dye according to the manufactureŕs recommendation (C10337; Invitrogen™, USA). To detect 53BP1, FANCD2, γ-H2AX and Cenp-C (also in micronuclei) cells were fixed with 4% paraformaldehyde for 10 minutes at room temperature and permeabilized with 0.5% Triton X-100 in PBS. Following permeabilization, cells were blocked with 3% BSA in PBS for 1 hour at room temperature. After blocking, cells were incubated overnight at 4°C with the following primary antibodies (diluted 1:750 in 3% BSA in PBS): anti-53BP1 (H-300; Santa Cruz, USA; cat no sc-22760; RRID: AB_2256326), anti-FANCD2 (Novus Biologicals, USA; cat no NB100-182; RRID: AB_3149940), anti-Phospho H2A.X (Ser139; γ-H2AX) (JBW301; mouse;

Merck Millipore, Germany; cat no #05-636; RRID: AB_309864), anti-CENP-C (MBL International, USA; cat no PD030; RRID: AB_10693556). This was followed by incubation with Alexa Fluor®-conjugated secondary antibodies (1:1000; Alexa Fluor 488, Alexa Fluor 594 and Alexa Fluor 647; Thermo Fisher Scientific, USA; cat no A-11029, A-11076, A-11034, A-11012 and A-21236) for 1 hour at room temperature. DNA was counterstained with Hoechst 33342 (1:10,000; Thermo Fisher Scientific, USA), then mounted onto glass slides using VECTASHIELD® antifade mounting medium (Vector Laboratories, USA). Images were acquired using a Leica DMI6000B fluorescence microscope equipped with a DFC360 FX camera.

### Proximity ligation assay for detection of transcription-replication conflicts (TRCs)

S-phase cells were labeled with 20 μM EdU (Jena Biosciences, Germany) for 30 minutes. Cells were pre-extracted with CSK100 buffer (100 mM NaCl, 300 mM sucrose, 3 mM MgCl₂, 10 mM MOPS, 0.5% Triton X-100 in PBS) and washed once with PBS. Cells were then fixed with 4% paraformaldehyde for 15 min, washed once with PBS, permeabilized with 0.5% Triton X-100 for 10 min, and washed again. EdU incorporation was detected using the Click-iT™ EdU Cell Proliferation Kit for Imaging with Alexa Fluor™ 594 dye (C10339; Invitrogen™, USA). Cells were blocked in 5% BSA for 1 h at 25 °C and incubated overnight at 4 °C with primary antibodies diluted 1:2000 in 5% BSA in PBS): anti-RNAPIIpS2 (1:2000; H5; BioLegend, USA; cat no. 920204; RRID: AB_2616695) and anti-PCNA (1:2000; Abcam, UK; cat no. ab18197; RRID: AB_444313), followed by two PBS washes. PLA was performed using the Duolink DUO92014 kit (Sigma-Aldrich, Germany) following the manufacturer’s protocol with slight modifications. Briefly, cells were incubated for 1 h at 37 °C with PLA probes (DUO92001 and DUO92005; 1:10 dilution in Duolink antibody diluent), followed by two 5-min washes in wash buffer A. Ligase was added (1:70 in 1× ligation buffer) and cells were incubated for 30 min at 37 °C, followed by two 5-min washes in wash buffer A. For amplification, polymerase was added (1:80 in 1× amplification buffer) and cells were incubated for 100 min at 37 °C, followed by two 10-min washes in wash buffer B. For DNA staining, DAPI (5 µg/ml) in PBS was added for 90 min, followed by two PBS washes. Cells were imaged at 40× magnification on a spinning disc confocal microscope (Andor Dragonfly). PLA foci were quantified using ImageJ after nuclei segmentation on the DAPI channel.

### Detection of chromosomal breaks and chromosome number variability

Chromosomal breaks and chromosome number variability were evaluated from metaphase chromosome spreads to assess CIN. Chromosome number variability was determined in single-cell clones cultured for 30 generations. Chromosomal breaks were assessed either in single-cell clones or in synchronized cells. For synchronized cells, a double thymidine block was used, followed by treatments applied from early to mid S phase. To enrich for early mitotic cells, asynchronous cells were treated with 2 μM Dimethylenastron (DME; Sigma-Aldrich, Germany) for 4–6 hours, whereas synchronized cells were treated with DME in late G2. Cells were then harvested and incubated in a hypotonic solution (40% culture medium/60% H_2_O) at room temperature for 15 minutes. Following incubation, the cell pellet was fixed in ice-cold Carnoy’s fixative (75% methanol/25% glacial acetic acid) and resuspended in 100% glacial acetic acid. The resulting cell suspension was dropped onto pre-chilled coverslips to generate chromosome spreads. Slides were air-dried and stained with 8% Giemsa solution (Sigma-Aldrich, Germany). Chromosome spreads were analyzed for chromosomal breaks and chromosomal number variability using a Zeiss Axioscope FS microscope equipped with a Hamamatsu C4742-95 camera or Leica DMI6000B fluorescence microscope equipped with a DFC360 FX camera. 50 chromosome spreads were evaluated per condition.

### CEP-FISH

Chromosome-specific aneuploidy was assessed using Fluorescence *in situ* Hybridization (FISH) detecting human chromosomes 1, 2, and 7. Single cell clones, cultured for 30 generations, were seeded onto coverslips, fixed with ice-cold Carnoy’s reagent (75% methanol, 25% glacial acetic acid), and air-dried. Coverslips were immersed in 2× SSC buffer (0.3 M NaCl, 0.03 M sodium citrate) for two minutes, followed by sequential dehydration in 70%, 85%, and 100% ethanol for two minutes each. Dried coverslips were hybridized with satellite enumeration probes specific to chromosomes 1, 2, and 7 (CytoCell, UK; LPE001G, LPE002R, LPE007R) following the manufacturer’s protocol. Fluorescent images of at least 50 nuclei per single cell clone were captured using a Leica DMI6000B fluorescence microscope equipped with a DFC360 FX camera (Leica, Germany). Image deconvolution was performed using Leica LAS AF software, and nuclei displaying chromosome counts deviating from the modal were quantified.

### Bioinformatic analysis

The Structural Complexity Score (SCS), Numerical Complexity Scores (NCS) and weighted genome instability index (wGII), as surrogate measures for structural, numerical and overall chromosomal instability (CIN), respectively, were calculated following the methodology described in ^27^. Plots were generated using the ggplot2, ggrepel, and ggpubr packages in R. The BiomaRt package in R was used to select genes from chromosome18q ^51,52^. To visualize the dependence of genomic loci on chromosomal instability, DNA copy data from The Cancer Genome Atlas (TCGA) was split based on CIN measures, with low CIN defined as being in the under the 40^th^ percentile and high CIN being above the 60^th^ percentile, using SCS and NCS. Frequency of genomic amplifications and deletions were calculated using the GISTIC2 module available on GenePattern (genepattern.org). This analysis was done separately for TCGA colorectal (CRC) and pancreatic ductal adenocarcinoma (PDAC) tumors. To assess the correlation between CIN-associated alterations in mRNA expression and the occurrence of mutations in genes located on chromosome 18q, differentially expressed genes were identified separately in TCGA and CCLE datasets and for both the SCS and the NCS using the limma-voom pipeline for RNAseq data implemented in the edgeR package in R. The resulting log fold changes (logFCs) were normalized and the mean logFCs were calculated. The adjusted p-values were pooled using Fisher’s method. For each gene, the rate of mutation associated with SCS and NCS in TCGA CRC and PDAC tumors was estimated using a linear model. The standardized mutation rates from the two CIN scores were then averaged to obtain a comparative measure of chromosome 18q mutations associated with CIN. To evaluate the association of *SMAD4* copy number loss to CIN, GISTIC2 thresholded copy number data for TCGA CRC and PDAC tumors was used. Samples were defined as displaying a loss of *SMAD4* if the GISTIC2 thresholded copy number was less than 0 (indicating a copy number loss), while wild-type samples were defined by a copy number of 0 (no alteration). To compare CIN in these two groups, a Wilcoxon rank sum test was applied to assess differences in wGII scores. To determine whether the positive correlation between *SMAD4* and *ID3* observed *in vitro* is also present in clinical samples, CorrelationAnalyzeR, a publicly available database of human disease (cancer vs. normal) and tissue-specific genome-wide co-expression correlation patterns, was utilized. This platform integrates uniformly processed RNA sequencing data from the Gene Expression Omnibus (GEO) and the Sequence Read Archive (SRA), curated by ARCHS4, for its analyses ^53^. All correlation analyses were performed on CRC and PDAC tissues using CorrelationAnalyzeR (https://gccri.bishop-lab.uthscsa.edu/shiny/correlation-analyzer/) ^54^.

### Publicly available data sources

CCLE RNA-seq counts: DepMap CCLE 2019 release [https://depmap.org/portal/data_page; accessed on August 23rd, 2023)] ^55^.

TCGA copy number segments, somatic mutations and RNA-seq counts: NCI’s Genomic Data Commons (GDC) data portal [https://portal.gdc.cancer.gov/; accessed on April 18th, 2023] ^56^.

TCGA GISTIC2 copy numbers: Xena browser TCGA PANCAN data hub [https://xenabrowser.net/hub/; accessed on June 27th, 2024] ^57^.

### Gene expression analysis and functional enrichment

Differentially expressed genes associated with *SMAD4* expression were identified using RNA-seq data from the TCGA colorectal cancer cohort. Gene-level counts were analyzed using the limma-voom pipeline implemented in the edgeR package in R. A linear model was fitted for each gene with *SMAD4* expression and tumor subtype (COAD vs. READ) as covariates to account for potential batch effects. P-values were adjusted for multiple testing using the Benjamini–Hochberg method. Genes meeting the significance threshold were submitted for Gene Ontology (GO) analysis using DAVID ^58^. Representative bubble charts were generated in R using ggplot2.

### Statistical analyses and graphics

Statistical analyses were performed using GraphPad Prism (version 10). Comparisons between two groups were performed using unpaired two-tailed Student’s t-tests (assuming unequal variances). Data are presented as mean ± standard deviation (SD). *P*-values are represented as follows: p ≥ 0.05p (ns), < 0.05 (*), p < 0.01 (**), p < 0.001 (***), and p < 0.0001 (****). Specific statistical tests used for other analyses are described in the corresponding figure legends. Elements in graphics in Fig. 3c, 3h and 6 were created using BioRender (https://BioRender.com)

## Data availability

All data, code, and materials used in the analysis will be available in some form to any researcher for purposes of reproducing or extending the analysis. The RNA sequencing data are available at GEO (accession number: GSE292334). Some materials will require an assignment of a material transfer agreement (MTAs). All data are available in the main text or the supplementary materials.

## Acknowledgments

The work was supported by grants from the Deutsche Forschungsgemeinschaft FOR2800 (H.B., M.K.) and KFO5002 (H.B., G.S.) and by the China Scholarship Council (CSC), File No. 202206220045 (S.Y.). The NGS-Sequencing instrument, the HiSeq4000 was funded by the Deutsche Forschungsgemeinschaft (DFG, German Research Foundation: INST 186/1217-1 FUGG). We thank Ali Bakr, Peter Schmezer, Linda Wordeman, Aristidis Moustakas and Zuzana Storchova for providing materials, Randolf Bodenstein and Magdalena Hennecke for technical help and Xiaoxiao Zhang for supporting bioinformatic analyses. We are grateful to Dana Branzei, Elisabeth Hessmann and members of the Bastians lab for comments on the manuscript.

## Author contributions

Conceptualization: HB, AP; Methodology: AP, GJ, PJ, SK, SY, IT, NB, MS, GS, SH, SH, MK, HB; Investigation: AP, GJ, PJ, SK, SY, IT, NB, MS, SH; Visualization: AP, GJ, PJ, SK, SY, IT, NB, MS, GS, SH, MK, HB; Funding acquisition: HB, GS, SH, MK; Project administration: HB; Writing: HB, AP; Review & editing: AP, GJ, SY, PJ, SK, MS, GS, SH, MK, HB

## Competing interests

Authors declare that they have no competing interests.

## Materials & correspondence

Requests for materials should be directed to the corresponding author Holger Bastians (email: holger.bastians@uni-goettingen)

## Notes

### Competing Interest Statement

The authors have declared no competing interest.

