## Supplemental Figures and Tables for "*SMAD4* loss drives cancer chromosomal instability through transcription-replication conflict-induced replication stress"

[1]University Medical Center Göttingen (UMG), Department of Molecular Oncology, Section for Cellular Oncology, Göttingen Center for Molecular Biosciences (GZMB), Göttingen, Germany. [2] University of Koblenz, Institute of Informatics, Koblenz, Germany. [3] University Medical Center Göttingen (UMG), Department of Pathology, NGS- Integrative Genomics Core Unit (NIG), Göttingen, Germany. [4] Helmholtz Center Munich, Institute of Epigenetics and Stem Cells (IES), Munich, 81377, Germany. [5] Clinical Research Unit 5002 (CRU5002), [6] Research Unit 2800 (FOR2800).

### Supplementary Figures and Figure Legends

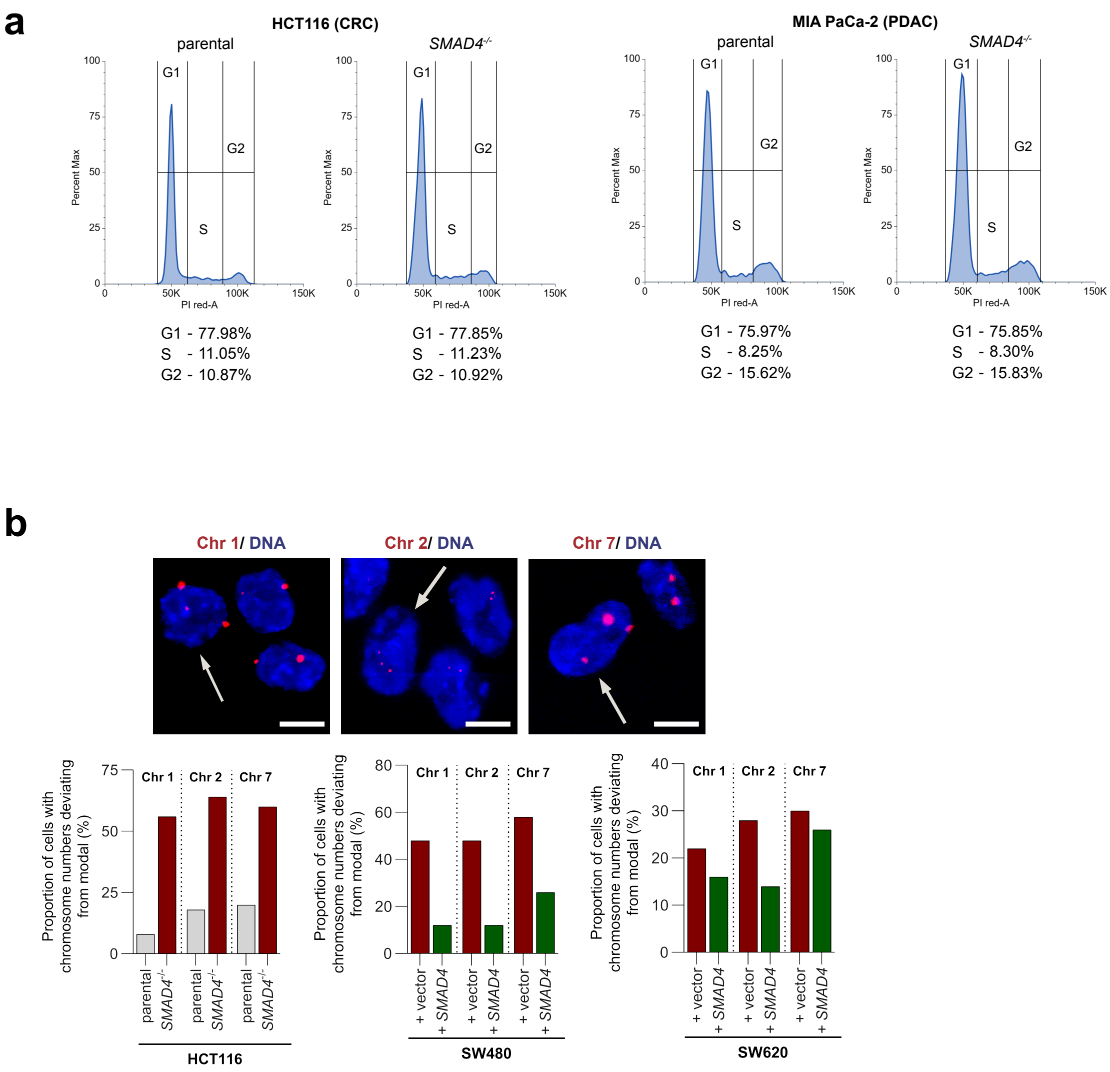


**Supplementary Figure 1: Loss of *SMAD4* does not affect cell cycle progression, but induces aneuploidy. a** Representative FACS profiles of CRC (HCT116) and PDAC (MIA PaCa-2) cells with or without *SMAD4* knockout. **b** Detection of aneuploidy using CEP-FISH in single cell clones derived from HCT116 cells with or without *SMAD4* knockout and from SW480 and SW620 cells with or without *SMAD4* re-expression. CEP-FISH analysis was used to detect chromosomes 1, 2 and 7 and the proportion of cells with CEP signals deviating from modal were determined (n=50 cells per condition). Example microscopy images for detection of chromosomes 1, 2 and 7 are given (scale bar, 10 µm).

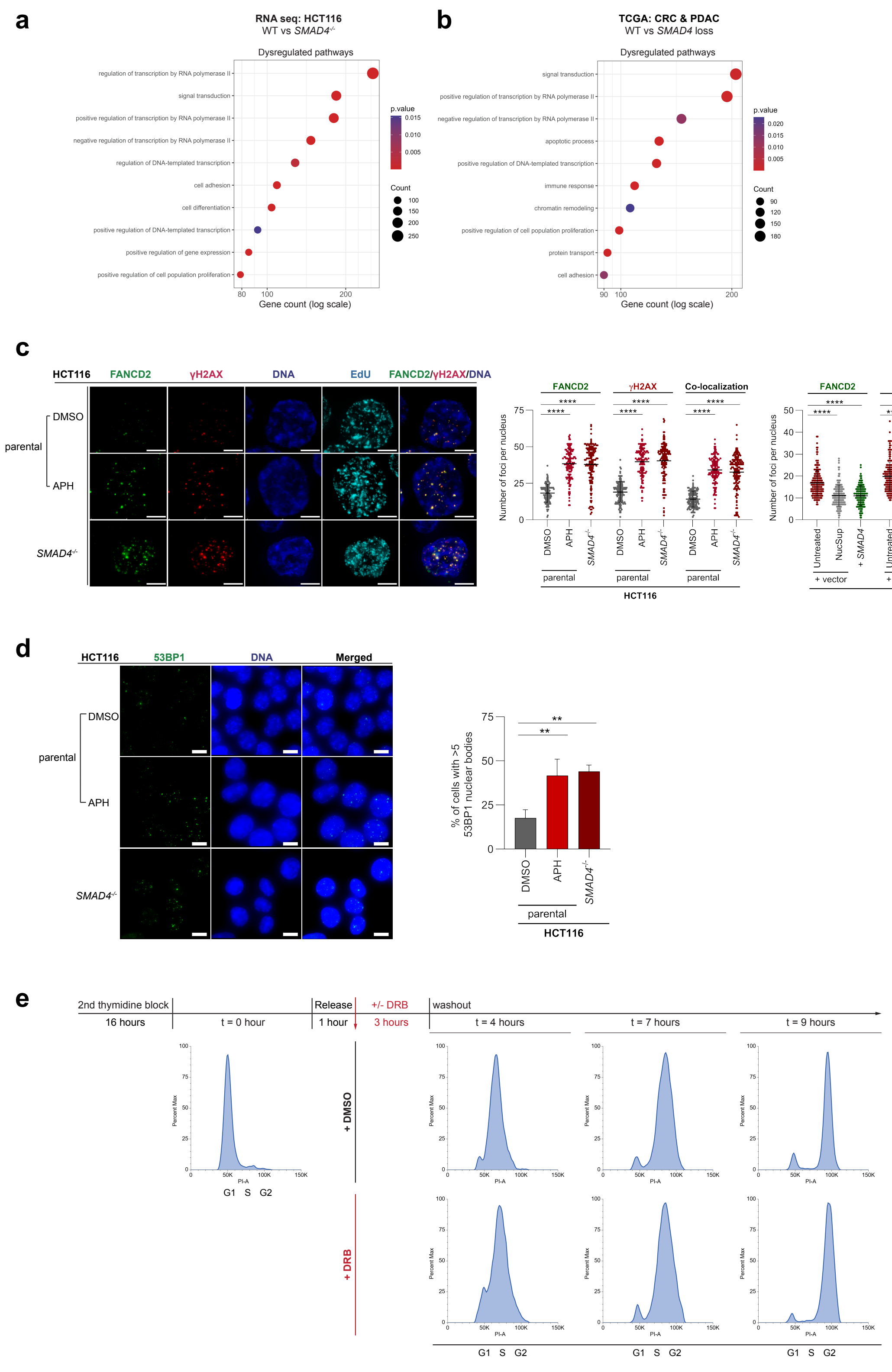
a
RNA seq: HCT116
WT vs SMAD4-/-
Dysregulated pathways
regulation of transcription by RNA polymerase II
signal transduction
positive regulation of transcription by RNA polymerase II
negative regulation of transcription by RNA polymerase II
regulation of DNA-templated transcription
cell adhesion
cell differentiation
positive regulation of DNA-templated transcription
positive regulation of gene expression
positive regulation of cell population proliferation
p.value
Count
Gene count (log scale)
b
TCGA: CRC & PDAC
WT vs SMAD4 loss
Dysregulated pathways
signal transduction
positive regulation of transcription by RNA polymerase II
negative regulation of transcription by RNA polymerase II
apoptotic process
positive regulation of DNA-templated transcription
immune response
chromatin remodeling
positive regulation of cell population proliferation
protein transport
cell adhesion
p.value
Count
Gene count (log scale)
c
HCT116
FANCD2
γH2AX
DNA
EdU
FANCD2/γH2AX/DNA
parental
DMSO
APH
SMAD4-/-
Number of foci per nucleus
Co-localization
****
Untreated
NucSup
+ SMAD4
+ vector
HCT116
SW480
d
HCT116
53BP1
DNA
Merged
% of cells with >5 53BP1 nuclear bodies
**
e
2nd thymidine block
16 hours
t = 0 hour
Release
1 hour
+/- DRB
3 hours
washout
t = 4 hours
t = 7 hours
t = 9 hours
+ DMSO
+ DRB
Percent Max
PI-A
G1 S G2

**Supplementary Figure 2: *SMAD4* loss dysregulates RNA polymerase II–regulated transcription and induces replication stress. a** Gene ontology analysis of RNA sequencing data from HCT116 cells with or without *SMAD4* knockout, revealing strong enrichment of pathways associated with RNA polymerase II-regulated transcription. **b** Gene ontology analysis of RNA sequencing data comparing *SMAD4*-low and *SMAD4*-high CRC (n=551) and PDAC (n=149) tumor samples from TCGA datasets, indicating perturbation of RNA polymerase II-regulated transcription. **c** Detection and quantification of FANCD2 and γ-H2AX nuclear foci and their colocalization during S phase by immunofluorescence microscopy in HCT116 cells treated with 100 nM aphidicolin to induce mild replication stress or upon *SMAD4* knockout or in *SMAD4*-deficient SW480 cells with or without *SMAD4* re-expression. SW480 cells were also supplemented with deoxynucleosides (NucSup) to alleviate replication stress. S phase cells were labelled by EdU incorporation and FANCD2, γ-H2AX and their colocalization were quantified. Example images are given (FANCD2, green; γ-H2AX, red; DNA, blue, EdU, cyan; scale bar, 5 µm). Bar graphs show mean ± SD (t-test, n=150-200 Edu-positive cells from 3 experiments). **d** Detection and quantification of 53BP1 nuclear bodies in HCT116 cells treated with 100 nM aphidicolin (APH) or after knockout of *SMAD4*. Representative microscopy images are given (53BP1, green; DNA, blue; scale bar, 10 µm). Bar graphs show the proportion of cells showing >5 53BP1 nuclear bodies (n= 300 cells from 3 experiments, mean ± SD, t-test). **e** Experimental outline and representative FACS profiles of HCT116 cells synchronized in S-phase and transiently treated with DRB to inhibit transcription elongation followed by washout of the drug.

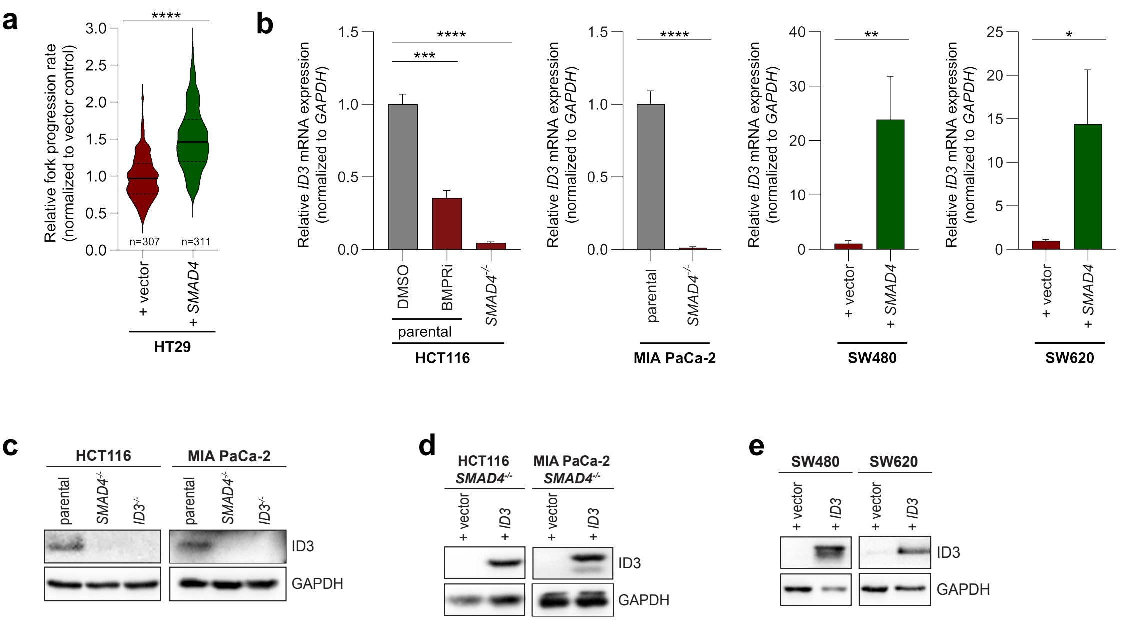


**Supplementary Figure 3: Identification of *ID3* as a transcriptional target of BMP-*SMAD4* signaling. a** Replication stress measurement by DNA combing in HT29 cancer cells with or without *SMAD4* re-expression (n=307-311 DNA fibers, mean ± SD, t-test). HT29 cells were used for RNA sequencing depicted in Fig. 4a. **b** Determination of relative *ID3* mRNA expression by qPCR analysis in HCT116 (CRC) and MIA PaCa-2 (PDAC) cells with or without *SMAD4* knockout or treated with 1 µM BMP receptor inhibitor and in SW480 and SW620 cells with or without *SMAD4* re-expression (mean ± SD, n=3 independent experiments, t-test). **c** Representative western blots showing loss of ID3 protein levels in HCT116 and MIA PaCa-2 cells with or without *SMAD4* or *ID3* knockout. **d** Representative western blots showing re-expression of *ID3* in HCT116- and MIA PaCa-2-*SMAD4*-knockout cells. **e** Representative western blots showing *ID3* re-expression in single cell clones derived from *SMAD4*-deficient SW480 and SW620 cells.

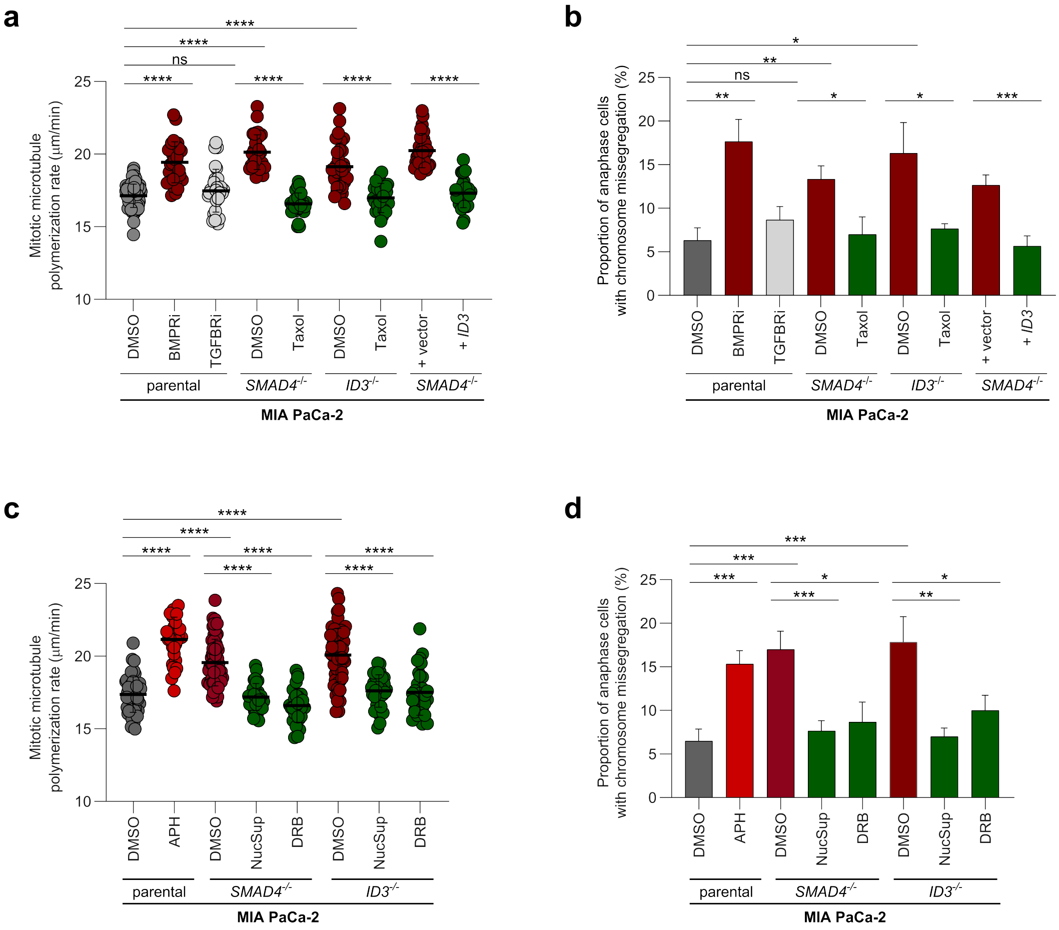


**Supplementary Figure 4: Inhibition of BMP/SMAD4/ID3 signaling causes increased spindle microtubule growth rates in mitosis and chromosome missegregation in PDAC cells. a** Measurements of mitotic microtubule growth rates in *SMAD4*-proficient MIA PaCa-2 cells treated with or without BMP or TGFβ receptor inhibitors, and in MIA PaCa-2-derived *SMAD4*- or *ID3*-knockout cells, with or without *ID3* re-expression. Cells were treated with 0.2 nM Taxol to suppress abnormally increased microtubule growth rates (mean ± SD, t-test, n=30 cells from 3 experiments with total of 600 microtubules measured). **b** Chromosome missegregation in MIA PaCa-2 according to the treatments used on (a) (mean ± SD, t-test, n=300 anaphase cells from 3 experiments). **c** Measurements of mitotic microtubule growth rates in MIA PaCa-2 cells with or without *SMAD4*- or *ID3*- knockout and treated with or without deoxynucleosides (NucSup) or DRB. Parental cells were treated with 100 nM aphidicolin (APH) to induce replication stress (mean ± SD, t-test, n=30 cells from 3 experiments with total of 600 microtubules measured). **d** Chromosome missegregation in MIA PaCa-2 cells according to the treatments as in (c) (mean ± SD, t-test, n=300 anaphase cells from 3 experiments).

### Supplementary Tables

**Supplementary Table 1:** Sequences of guide RNAs used for CRISPR-Cas9-mediated targeting of *SMAD4* and *ID3*

| **Name** | **Target** | **Strand** | **Sequence** | **PAM** |
|---|---|---|---|---|
| Hs.Cas9.SMAD4.1.AB | *SMAD4* | + | TACGAACGAGTTGTATCACC | TGG |
| Hs.Cas9.SMAD4.1.AC | *SMAD4* | - | TCTGCAACAGTCCTTCACTA | TGG |
| CD.Cas9.RPMB8615.AI | *ID3* | - | TCGTTGGAGATGACAAGTTC | CGG |
| Hs.Cas9.ID3.1.AA | *ID3* | + | TGGCTAAGCTGAGTGCCTCT | CGG |

**Supplementary Table 2:** Sequences of oligonucleotides used for target gene amplification via PCR

| **Name** | **Target** | **Sequence** |
|---|---|---|
| *ID3*-forward | CD.Cas9.RPMB8615.AI; Hs.Cas9.ID3.1.AA | 5’- TTGCTGGACGACATGAACCA - 3’ |
| *ID3*-reverse | | 5’- ACTCCAGGACTTGCCGTTTA -3’ |
| *SMAD4*-forward | Hs.Cas9.SMAD4.1.AB | 5’- TGATCTATGCCCGTCTCTGGA - 3’ |
| *SMAD4*-reverse | Hs.Cas9.SMAD4.1.AB | 5’- TAATGTTACTGCCTGCCGCTC - 3’ |
| *SMAD4*-forward | Hs.Cas9.SMAD4.1.AC | 5’- GCCAGACGTACAGTGGTGTT - 3’ |
| *SMAD4*-reverse | Hs.Cas9.SMAD4.1.AC | 5’- CAGTCCAGGTGGTAGTGCTG - 3’ |

**Supplementary Table 3:** Sequences of oligonucleotides used for qRT-PCR analysis

| **Name** | **Target** | **Sequence** |
|---|---|---|
| *ID3*-forward | *ID3* | 5'- TCATCTCCAACGACAAAAGG -3' |
| *ID3*-reverse | *ID3* | 5'- ACCAGGTTTAGTCTCCAGGAA -3' |
| *GAPDH*-forward | *GAPDH* | 5'- GAAGGTCGGAGTCAACGGATT- 3' |
| *GAPDH* -reverse | *GAPDH* | 5'- CGCTCCTGGAAGATGGTGAT -3' |
